# Disentangling the effects of eutrophication and natural variability on macrobenthic communities across French coastal lagoons

**DOI:** 10.1101/2022.08.18.504439

**Authors:** Auriane G. Jones, Gauthier Schaal, Aurélien Boyé, Marie Creemers, Valérie Derolez, Nicolas Desroy, Annie Fiandrino, Théophile L. Mouton, Monique Simier, Niamh Smith, Vincent Ouisse

## Abstract

Coastal lagoons are transitional ecosystems that host a unique diversity of species and support many ecosystem services. Owing to their position at the interface between land and sea, they are also subject to increasing human impacts, which alter their ecological functioning. Because coastal lagoons are naturally highly variable in their environmental conditions, disentangling the effects of anthropogenic disturbances like eutrophication from those of natural variability is a challenging, yet necessary issue to address. Here, we analyze a dataset composed of macrobenthic invertebrate abundances and environmental variables (hydro-morphology, water, sediment and macrophytes) gathered across 29 Mediterranean coastal lagoons located in France, to characterize the main drivers of community composition and structure. Using correlograms, linear models and variance partitioning, we found that lagoon hydro-morphology (connection to the sea and lagoon surface), which affects the level of environmental variability (salinity and temperature), as well as lagoon-scale benthic habitat diversity (using macrophyte morphotypes) seemed to regulate macrofauna distribution, while eutrophication and associated stressors like low dissolved oxygen, acted upon the existing communities, mainly by reducing species richness and diversity. Furthermore, M-AMBI, a multivariate index composed of species richness, Shannon diversity and AMBI (AZTI’s Marine Biotic Index) and currently used to evaluate the ecological state of French coastal lagoons, was more sensitive to eutrophication (18%) than to natural variability (9%), with nonetheless 49% of its variability explained jointly by both. To improve the robustness of benthic indicators like M-AMBI and increase the effectiveness of lagoon benthic habitat management, we call for a revision of the ecological groups at the base of the AMBI index and of the current lagoon typology which could be inspired by the lagoon-sea connection levels used in this study.

## Introduction

Coastal lagoons are inland, shallow and transitional water bodies, usually oriented parallel to the coast, separated from the sea by a barrier, connected to the sea by one or more restricted inlets which remain open at least intermittently (Kjerfve, 1994). These ecosystems represent 13% of the world’s coastline (Barnes, 1980) and provide essential ecosystem services, especially through nutrient regulation and sequestration as well as climate regulation (Kermagoret et al., 2019; Levin et al., 2001). These ecosystems are characterized by strong salinity variations at different temporal scales, ranging from fortnights to seasons and decades (Pérez-Ruzafa et al., 2007a), that are mainly governed by short-term freshwater inflow, seasonal temperature variations, the degree of connection of each water mass with the sea and decadal climatic variations (rainfall and temperatures) (Barbone and Basset, 2010; De Casabianca, 1996; Zaldívar et al., 2008). Oxygen saturation naturally fluctuates in lagoons, due to water temperature and re-aeration processes controlled by wind conditions (Chapelle et al., 2001; Derolez et al., 2020a). Oxygen dynamics are further affected by primary production, which is mainly governed, in oligo and mesotrophic lagoons by seasonal benthic macrophyte growth and decay cycles, and in hyper and eutrophic lagoons by pelagic phytoplankton dynamics (Zaldívar et al., 2008). Overall, coastal lagoons are naturally highly dynamic systems in terms of environmental factors, representing a form of natural environmental stress for any organism living in these systems like fish and benthic invertebrates (Pérez-Ruzafa et al., 2007b; Zaldívar et al., 2008). Consequently, lagoon benthic communities, just like estuarine ones, generally present low species richness and high abundances of tolerant species, characteristics typical of disturbed or stressed coastal ecosystems, giving rise to the “transitional waters quality paradox” (Zaldívar et al., 2008), a generalization of the “Estuarine quality paradox” (Elliott and Quintino, 2007). These paradoxes highlight the problems faced by transitional waters concerning the development of methodologies permitting the evaluation of their ecological quality status, because it results from both anthropogenic pressures and natural characteristics.

Coastal lagoons have also been intensely used and modified by humans for food (fishing and aquaculture), transportation (canals, ports and inlet modification), livelihood (land reclamation for construction) and recreation (fishing, sailing, swimming) (Lasserre, 1979; Pérez-Ruzafa et al., 2011a). The position of coastal lagoons at the interface between land and sea further exposes them to human land-derived pollution such as nitrogen, phosphorous, pesticide, or heavy metal pollution (Caliceti et al., 2002; Derolez et al., 2019; Souchu et al., 2010). The oldest and most studied is nitrogen and phosphorous pollution linked to agriculture and urban development (Bendoricchio et al., 1993; Kjerfve, 1994; Lacoste et al., 2023; Tournoud et al., 2006), which can lead to eutrophication of coastal lagoons, a phenomenon associated to an increase in phytoplankton biomass (Cloern, 2001; Zaldívar et al., 2008), a replacement of long-lived submerged macrophytes like marine plants, by emergent free-living benthic macroalgae and even to the disappearance of all benthic macrophytes (Le Fur et al., 2019, 2018). The high light availability linked to shallow depths and the naturally high nutrient concentration of water and sediments (*i.e.* coastal lagoons can be considered as “naturally eutrophic” (De Casabianca, 1996; Zaldívar et al., 2008)) promote the development of structurally complex benthic macrophytes, which are characteristic features of coastal lagoons. Macrophyte presence, cover and composition create a highly heterogeneous aquatic landscape (Arocena, 2007; Gamito et al., 2005), which is also determined by the sediment nature (*i.e.* hard bottom, gravel, sand, mud) and organic matter content (Menu et al., 2019).

Benthic macroinvertebrates are a key biotic compartment of coastal lagoons, through their role in water filtration, food web dynamics, recycling of organic matter and nutrients and sediment bioturbation (Barnes, 1980; Gamito et al., 2005; Levin et al., 2001; Zaldívar et al., 2008). These benthic communities are reported to be shaped mainly by natural environmental variability (*i.e.* environmental filtering) and passive diffusion (mainly as planktonic larvae) (Basset et al., 2007). At the intra-lagoon scale, the spatial distribution of macrobenthic invertebrates is tightly linked to the renewal of marine-originating molecules, which is directly linked to the connection level with the open sea, a result used to define the notion of confinement (Guelorget and Perthuisot, 1983). Superimposed onto the previously mentioned “natural environmental stress” (*i.e.* strong fluctuations of abiotic parameters like salinity and temperature) are a number of human related disturbances like anthropogenic eutrophication, which can further impact lagoonal organisms (Elliott and Quintino, 2007; Rossi et al., 2006). Most benthic macroinvertebrates are exposed to multiple human-induced disturbances as they generally live for several years. Furthermore, these organisms tend to present, as adults, a low dispersal potential (*i.e.* below a few kilometers), meaning that faced with a disturbance, each individual will either resist, become accustomed or die, with overall very little chance of escaping the disturbance, especially if it affects the entire lagoon (Cognetti and Maltagliati, 2000). These characteristics make benthic macroinvertebrates good theoretical indicators of the ecological health status of coastal lagoons and since 2000, this biological compartment is officially included in the assessment of European transitional water bodies carried out within the Water Framework Directive (WFD) (European Commission, 2018). However, although they are in theory good ecological indicators, benthic macroinvertebrates are also influenced by “natural environmental stress”, hence it appears crucial to disentangle the effect of natural environmental variability from the effect of variability linked to human disturbances on this compartment in hyper-variable ecosystems like coastal lagoons (Pitacco et al., 2019). Most of the published work on Mediterranean coastal lagoon macroinvertebrates have either been descriptive studies focusing on one or a few lagoons (Amanieu et al., 1977; Arocena, 2007; Bachelet et al., 2000; Carlier et al., 2007; Guelorget et al., 1994; Guelorget and Michel, 1979; Magni et al., 2022; Pérez-Ruzafa et al., 2007a; Rossi et al., 2006), theoretical oriented studies testing ecological hypotheses like environmental filtering (Basset et al., 2007) or species-area relations (Sabetta et al., 2007) or very applied studies related to the WFD and focusing on lagoon typology definition, indicator development or indicator reference values (Barbone et al., 2012; Basset et al., 2013a; Reizopoulou et al., 2014). Disentangling the respective effect of natural vs human-induced disturbances on lagoon communities requires a statistical power which can be reached by using extensive datasets, spanning a variety of ecological conditions (*i.e.* several lagoon systems) and a thorough characterization of both sources of disturbance. In this study, we make use of the full potential of large-scale environmental surveys of the various physical and biological compartments (water column, sediments, macrophytes and benthic macroinvertebrates) of French Mediterranean coastal lagoons: *(i)* to disentangle the effects of an anthropogenic disturbance, here eutrophication, from those of natural environmental variability on benthic macroinvertebrates and *(ii)* to understand the links between the different environmental variables characterizing coastal lagoons, that affect benthic macroinvertebrates. This study, based on a dataset including 29 French Mediterranean coastal lagoons (ranging from oligo- to eu-haline, with diverse environmental and anthropic settings), focuses on multiple levels of biological complexity; first the broad and complex community level, then simple diversity indices like species richness and Shannon diversity index and finally, a multimetric and WFD-compliant index.

## Methods

### Study sites

The French Mediterranean coast features a great diversity of lagoons characterized by different hydro-climatic contexts (*e.g.* temperature, wind, rainfall). Some of these lagoons are under the influence of large watersheds (up to 1 500 km²) while others are connected to more restricted basins that may be 300 times smaller (*ca.* 5 km²). These lagoons also present variable connections to the sea, with both natural vs man-made channels, and continuous vs temporary channels. Human land-use in this region is particularly high, associated to both urban and agricultural development (Derolez et al., 2019). Most lagoons in continental France are located near large cities like Narbonne or Montpellier and/or in areas intensively cultivated for wine, vegetables and cereals (CORINE Land Cover, 2018) whereas the Corsican lagoons of Palo, Urbino and Diana are relatively preserved from direct urban pressures. Differently, the Corsican lagoon of Biguglia is located just south of the urban hub of Bastia and in a growing agricultural plain (CORINE Land Cover, 2018; Pasqualini et al., 2017). Overall, French Mediterranean lagoons face various degrees of anthropogenic pressures.

A total of 29 nanotidal (tide < 0.5 m) coastal lagoons located in the Gulf of Lion (north-west Mediterranean Sea) and along the east coast of Corsica were selected for this study and sampled across 41 stations (between one and three in each lagoon) (Figure 1). The sampling was carried out once a time in 2006 or 2009 (depending on the station) during the “Réseau de Suivi Lagunaire” and WFD campaigns. Infaunal benthic macroinvertebrates were sampled in May, except four lagoons sampled at the end of April (La Marette, Ponant, Berre and Vaine) and one lagoon at the end of June (Bolmon). The studied lagoons encompass a wide range of hydro-morphological characteristics in terms of surface, depth and connection to the sea (Table 1) with the lagoon-sea connection level expressed as weak, moderate or strong depending on the differences in salinity variations between the lagoon and a neighboring marine station (Derolez et al., 2012). Lagoon surfaces vary from 1 km^2^ (Campignol) to 133 km^2^ (Berre) (Menu et al., 2019) and station depth is comprised between 0.1 m (La Palme south station) and 9.5 m (Urbino) with most stations less than 3 m deep (Table 1).

**Figure 1.**
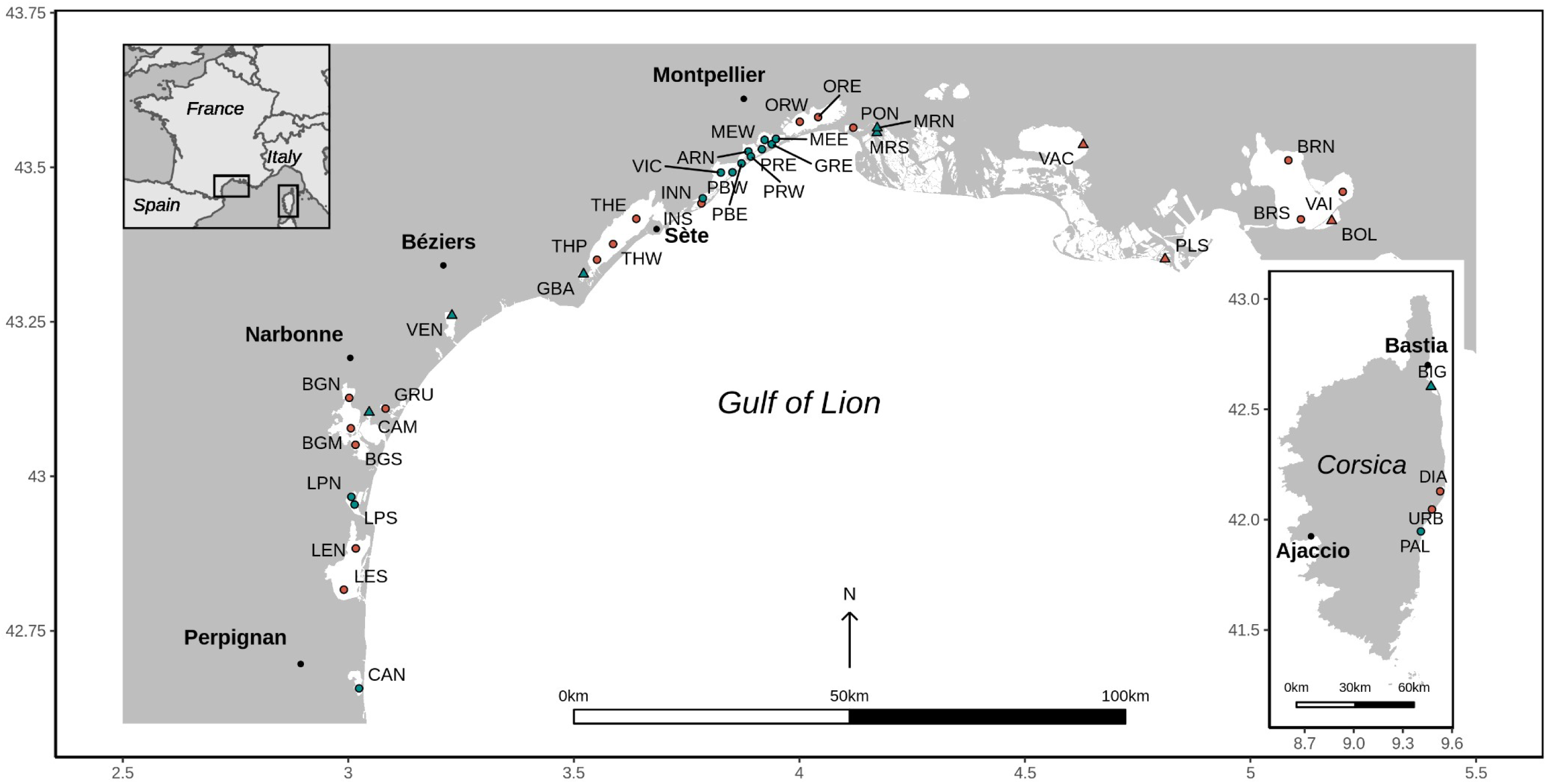
Map of the 41 stations sampled in the 29 lagoons located along the French Mediterranean coast (Gulf of Lion and Corsica). Each station’s salinity type (according to *their median salinity and Venice classification (Anonymous, 1958) in* Le Fur et al. (2018)) and group membership (K-means partitioning) based on the benthic macrofauna Hellinger-transformed mean abundances are indicated using a symbol (triangle: oligo- and meso-haline and circle: poly- and eu-haline) and a color (red: group A and blue: group B), respectively (see section *Macrobenthic community structure* for the explanation on this analysis). See Table 1 for the full lagoon names and corresponding stations.

**Table 1.**
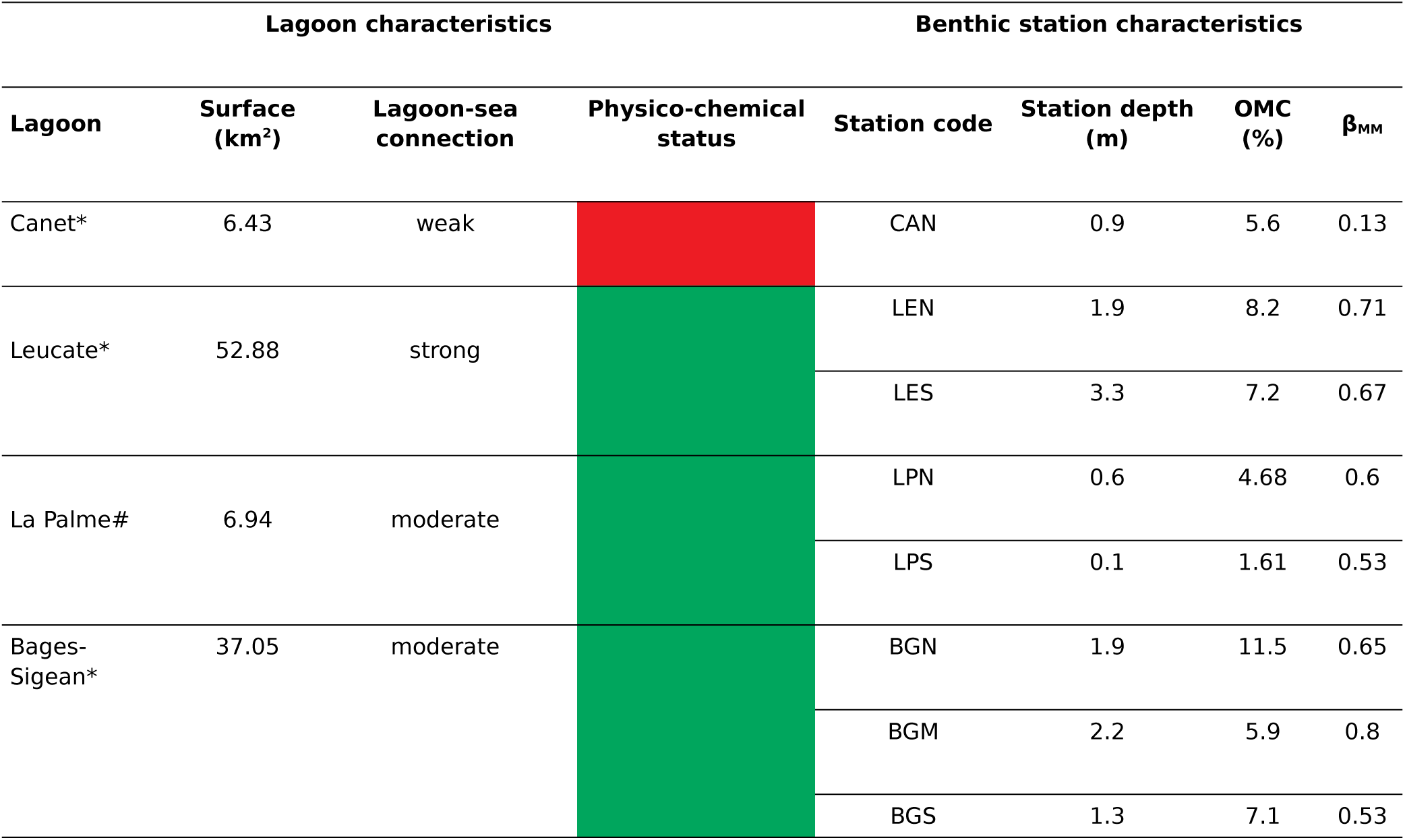
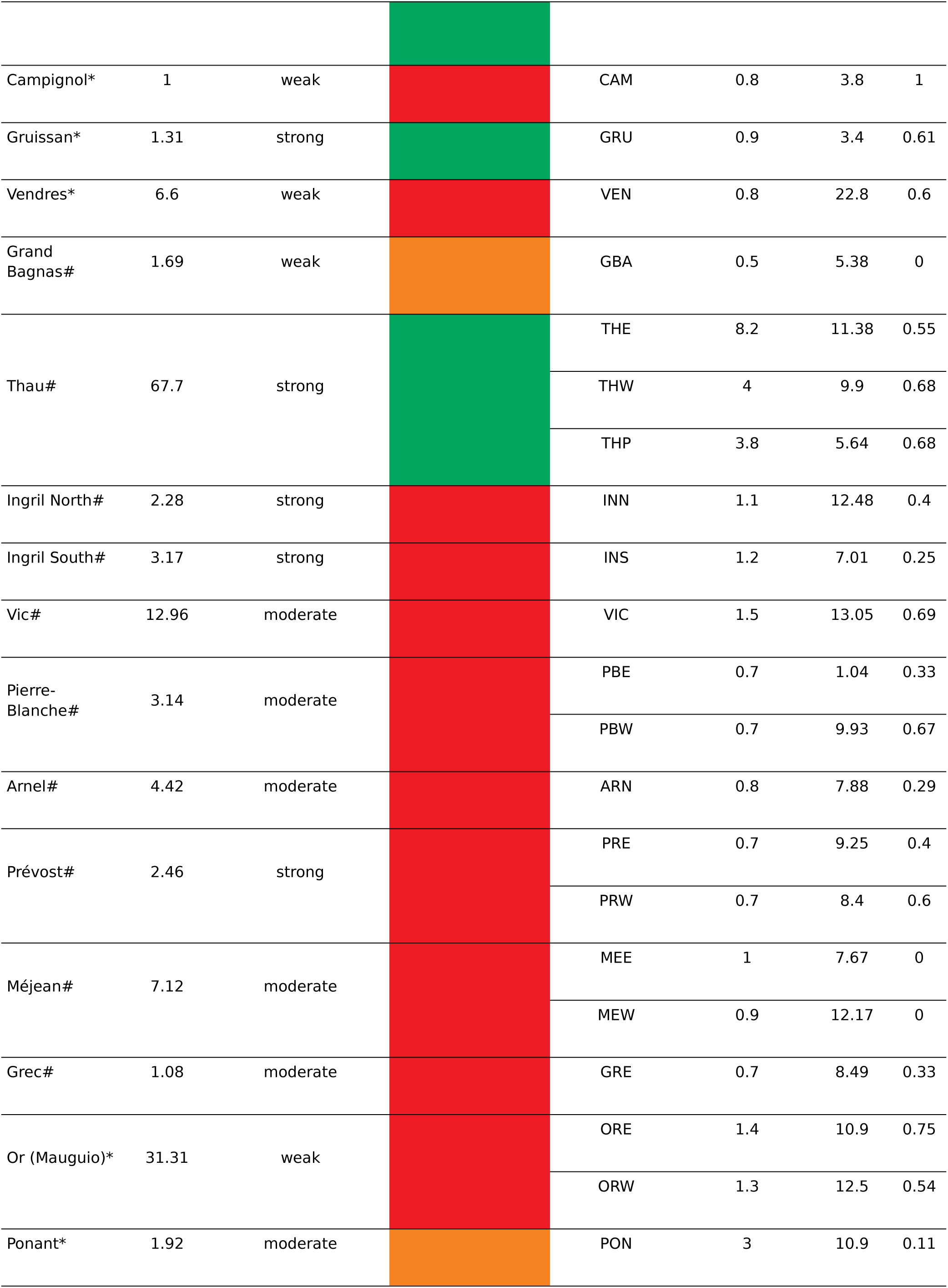
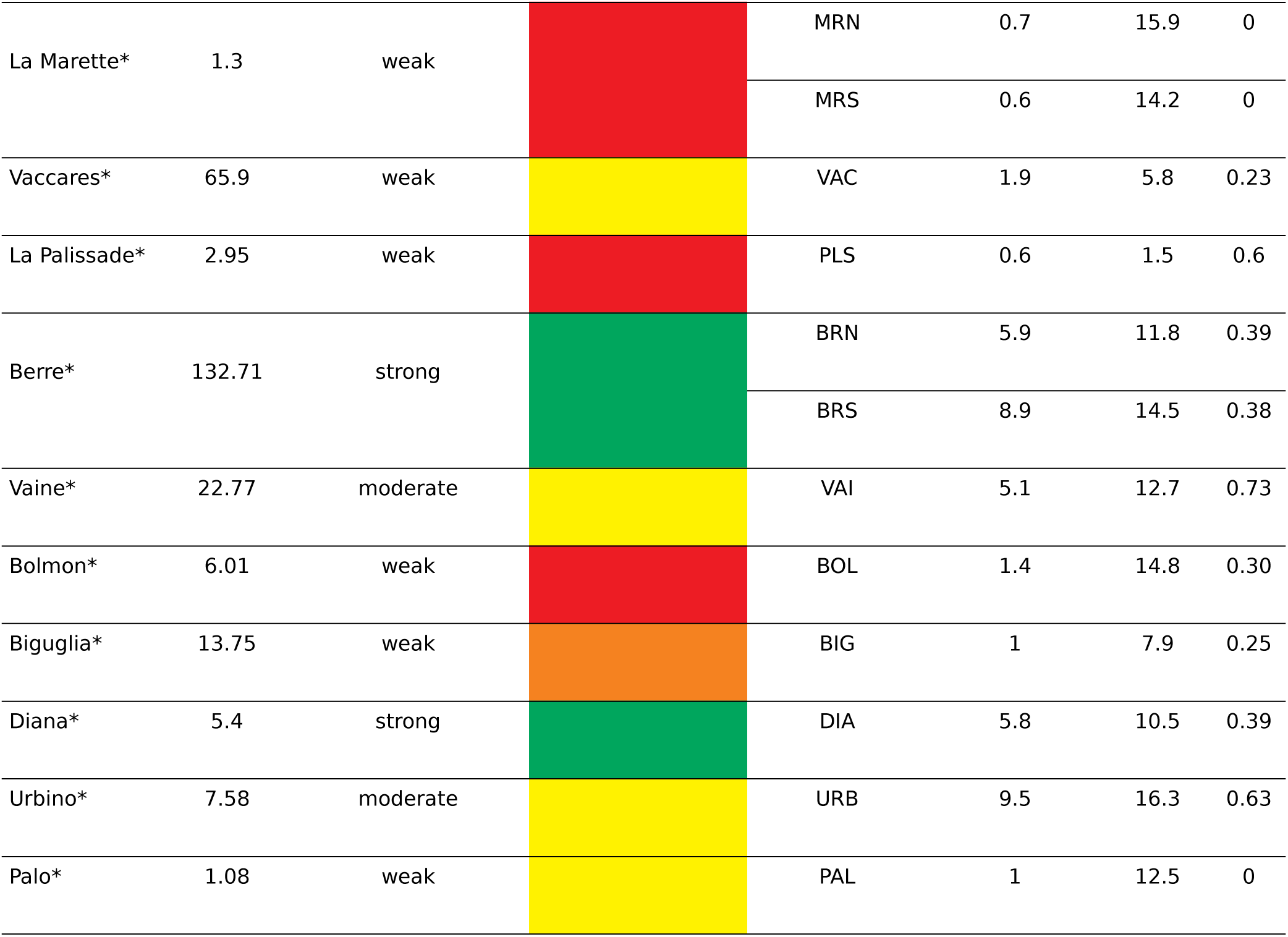
Hydro-morphological characteristics (surface, depth and connection level to the sea) and water physico-chemical status of the 29 lagoons as evaluated by the water framework directive (green = good, yellow = moderate, orange = bad, red = poor). The physico-chemical status is based on concentrations of total and dissolved inorganic forms of nutrients, turbidity and oxygen saturation during the summer months of the 2004-2009 period (Andral and Sargian, 2010a; Andral and Sargian, 2010b). Depth, sediment organic matter content (OMC, % of dry sediment weight), beta diversity of macrophyte morphotypes (β_MM_, no unit – see section *Estimating the macrophyte morphotype beta diversity*) of the 41 stations sampled for benthic macrofauna either in 2006 (#) or in 2009 (*) is also indicated. For stations sampled in the same lagoon, each one is labeled using a 3 letter code with the last letter indicating the geographical location of the station in the lagoon (North (N), Middle (M), South (S), East (E), West (W), except THP which indicates the station’s local name Pisse-Saumes). The lagoons are listed from west to east along the Gulf of Lion (see Figure 1 for their geographical location).

The different hydro-morphological characteristics of the studied lagoons lead to a great diversity of salinity regimes with 8 oligo- or meso-haline lagoons and 21 poly- or eu-haline lagoons (according to Le Fur et al., (2018), Figure 1) and to large discrepancies between lagoons in terms of annual variability of temperature, salinity and oxygen saturation. Overall, the physico-chemical status of the studied lagoons between 2004 and 2009 was evaluated according to the WFD methodology, as poor (15 lagoons), bad (3 lagoons), moderate (4 lagoons) or good (7 lagoons) (Table 1). This evaluation takes into account the concentrations of total and dissolved inorganic forms of nutrients, turbidity and oxygen saturation during the summer months of the 2004-2009 period and is conducted according to the WFD quality grid (Andral and Sargian, 2010a, 2010b).

### Sampling

#### Benthic macroinvertebrates

At each of the 41 stations, three replicates (*ca.* 10 m apart) of unvegetated sediment were sampled. Each replicate was made up of four pooled Ekman-Birge grabs, representing a total surface of 0.09 m^2^ per replicate and 0.27 m^2^ per station. Samples were sieved on a 1 mm mesh, fixed in a 5 to 7% formaldehyde/seawater solution and stained with Rose Bengal for latter identification. The sediments were then sorted in the laboratory and all organisms identified if possible to the species level, following the nomenclature of the European Register of Marine Species (ERMS). Taxa are indicated using WORMS reference (https://www.marinespecies.org/aphia.php?p=match). Prior to the data analyses, we removed meiofauna taxa (Ostracoda, Nematoda, Harpacticoida and Collembola) and organisms present in only one lagoon and determined at the order (Decapoda, Coleoptera, Nudibranchia), class (Insecta, Ascidiacea) or subclass (Oligochaeta) level. We kept Actiniaria, Nemertea and Polycladida as they were identified across multiple lagoons.

#### Sediment parameters

At each station, an additional grab sample was collected to evaluate sediment organic matter and grain-size distribution. Sediment samples were sieved on a 2 mm mesh to remove large macrophytes debris, and the filtrate was weighted and dried until stable weight (110°C for 24 to 48h) in the laboratory. Organic matter content (OMC, % of dry sediment weight) was calculated as the loss of weight after combustion for 12h at 450°C of the dry sediment fraction below 2 mm. Because of technical problems, no OMC was available for the stations ARN, LPN, LPS, PRW and VIC in 2006. To estimate these missing values, we used OMC values measured in spring of 2006 (ARN, PRW, VIC) or spring of 2007 (LPN and LPS) for the WFD sediment quality evaluation. For ARN, LPS, LPN and VIC, we estimated OMC by averaging over the values measured in the three or four stations closest to the macrofauna station (distance between 285 m and 707 m). For PRW, we used the OMC value measured at the closest station as this station was located only 114 m from the macrofauna station (Supplementary material 1 and 2). Since laboratory protocols changed between years (2006 and 2009) and between lagoons regarding the sediment particle sizes considered for the grain-size analysis, we chose not to consider grain-size parameters.

#### Water physico-chemical properties

Water column physico-chemical properties of the 29 lagoons are measured monthly during summer (from June to August or September) at least since 2003, following a standardized protocol implemented by the WFD (Derolez et al., 2019). Salinity (Sal), temperature (Temp) and oxygen saturation (O_2_ sat) were measured using a portable field probe (WTW LF 197, WTW-GmbH, Weilheim, Germany) and turbidity (Turb) was measured in the laboratory using an optical turbidimeter (2100N IS turbidimeter ISO 7027). Chlorophyll *a* concentration ([Chla]) was measured in 50 mL samples of water, first filtered on a Whatman GF/F filter (0.7 µm). Pigments were later extracted in the laboratory in a 90% acetone + distilled water solution during 24 hours (4°C, dark conditions after 10 minutes of centrifugation). The supernatant was then dosed using a spectrofluorometer following Neveux and Lantoine (1993). The full description of the protocol followed to measure [Chla], total nitrogen concentration [TN], ammonium concentration [NH_4_^+^], sum of nitrate plus nitrite concentrations [NO_x_], total phosphorus concentration [TP] and dissolved inorganic phosphorus concentration [DIP] is described in Bec et al. (2011) and Souchu et al. (2010). Only sub-surface data were available for all stations except the THW, THE, BRN, BRS stations located in the deep Thau and Berre lagoons for which sub-surface and bottom data were available. In these four cases, we chose to consider the bottom data.

#### Estimating the macrophyte morphotype beta diversity

Lagoons are characterized by a mosaic of benthic habitats, determined in particular by macrophyte diversity which can affect benthic macroinvertebrates (Arocena, 2007; Cardoso et al., 2004; Magni and Gravina, 2023; Ouisse et al., 2011) by acting as food (Jones et al., 2021; Quillien et al., 2016; Vizzini and Mazzola, 2006), refuge (Mancinelli and Rossi, 2001; Nordström and Booth, 2007; Ware et al., 2019) and/or means of transport (Norkko et al., 2000). Although sampling was carried out on locally unvegetated sediments to avoid biases linked to macrophytes and associated faunal assemblages, we postulate that the presence and composition of macrophyte assemblages within each lagoon can affect the composition of infaunal benthic communities. We developed an index (later beta diversity of macrophyte morphotypes, β_MM_) calculated as the average Bray-Curtis dissimilarity at the lagoon or sub-lagoon scale (when several macrofauna stations were sampled in one lagoon) based on a station by macrophyte morphotype matrix (presence/absence). Each macrophyte station is coded as bare sediment if the macrophyte cover is less than 25% or as one or several of seven macrophyte morphotypes (*i.e.* mushroom, feather, hair, ball, branched, sheet and meadow) based on the thallus morphology of the dominant or co-dominant macrophyte taxa. This index varies between 0 if all the considered stations are dominated or co-dominated by the same macrophyte morphotype(s) or if they all have bare sediment and 1 if all the considered stations are dominated or co-dominated by different macrophyte morphotypes (Appendices 1 and 2). We gathered the data used to calculate this index from the WFD macrophyte dataset (all information regarding sampling protocols are available in Le Fur et al., 2017).

### Data analyses

We extracted all the data used in this study from the French database Quadrige (Ifremer, http://quadrige.eaufrance.fr/). All the raw data used in this study (macrofauna, environmental data, GIS data) are available online along with all the scripts and code used to analyze the data (Jones et al., 2024). We performed all analyses using R version 3.6.3 (R Core Team, 2020) and the packages vegan (*v2.5-*6; Oksanen et al., 2019), missMDA (Josse and Husson, 2016), labdsv (*v2.0-*1; Roberts, 2019), olsrr (v0.5.3; Hebbali, 2020), MASS (Venables and Ripley, 2002), corrplot (*v0.84*; Wei and Simko, 2017) and rstatix (v0.6.0; Kassambara, 2020). We considered a significance level of 0.05 for all statistical tests.

#### Environmental variables

Water column parameters were measured in summer whereas macrofauna was sampled in spring which prevented us from relating these two datasets. Nonetheless, the presence and abundance of macroinvertebrates in a lagoon year y is influenced not only by the environmental conditions prevailing that year but also those prevailing the previous years especially for pluri-annual species. As such, we associated the macrofauna data collected year y to the mean and coefficient of variation (CV) of the water column parameters measured years y-2, y-1 and y, as done by Le Fur et al. (2019) (Table 2).

**Table 2.** Environmental parameters and their respective code used to characterize the water. For each water parameter, the mean and coefficient of variation (CV) were calculated between the years y-2 and y (with y the macrofauna sampling year) and the overall median, minimal and maximal values across the 41 stations are indicated. The station with the minimal and maximal value of each parameter is also indicated between brackets, as well as the transformation applied to each parameter. See Table 1 for the station codes.

| Parameter (unit) | Code | Measure | Median | Min | Max | Transf. |
| --- | --- | --- | --- | --- | --- | --- |
| Salinity (PSU) | Sal | mean | 32.39 | 2.83 (PLS) | 42.30 (PBW) | / |
|  |  | CV | 13.52 | 2.49 (DIA) | 82.24 (CAM) | log<sub>e</sub> |
| Oxygen saturation (%) | O<sub>2</sub> sat | mean | 100.88 | 56.61 (BRN, BRS) | 190.40 (BOL) | log<sub>e</sub> |
|  |  | CV | 27.38 | 0.82 (PLS) | 67.33 (PRE) | √ |
| Temperature (°C) | Temp | mean | 24.08 | 19.63 (BRN, BRS) | 27.09 (URB, BIG) | / |
|  |  | CV | 9.05 | 1.82 (CAN) | 22.81 (PON) | √ |
| Chlorophyll a concentration (µg.L<sup>-1</sup>) | [Chla] | mean | 9.43 | 0.70 (BGS) | 144.20 (MEW) | log<sub>e</sub> |
|  |  | CV | 71.04 | 25.42 (BOL) | 178.73 (BIG) | log<sub>e</sub> |
| Total nitrogen concentration (µmol.L<sup>-1</sup>) | [TN] | mean | 52.05 | 11.08 (DIA) | 266.03 (MEW) | log<sub>e</sub> |
|  |  | CV | 27.77 | 6.94 (VAC) | 89.47 (BIG) | log<sub>e</sub> |
| Ammonium concentration (µmol.L<sup>-1</sup>) | [NH<sub>4</sub><sup>+</sup>] | mean | 1.47 | 0.06 (PLS) | 31.62 (GRE) | log<sub>e</sub> |
|  |  | CV | 106.88 | 28.65 (CAN) | 257.93 (THE) | √ |
| Nitrate plus nitrite concentration (µmol.L<sup>-1</sup>) | [NO<sub>x</sub>] | mean | 0.37 | 0.06 (GBA) | 22.81 (CAM) | log<sub>e</sub> |
|  |  | CV | 94.87 | 43.30 (CAN) | 227.37 (URB) | log<sub>e</sub> |
| Total phosphorous concentration (µmol.L<sup>-1</sup>) | [TP] | mean | 2.51 | 0.64 (LEN) | 31.19 (CAN) | log<sub>e</sub> |
|  |  | CV | 36.75 | 8.91 (VAC) | 106.12 (ARN) | log<sub>e</sub> |
| Phosphate concentration (µmol.L<sup>-1</sup>) | [DIP] | mean | 0.24 | 0.02 (LES) | 14.53 (CAN) | log<sub>e</sub> |
|  |  | CV | 93.10 | 40.97 (THW) | 187.55 (INN) | log<sub>e</sub> |
| Turbidity (NTU) | Turb | mean | 5.20 | 1.14 (BGS) | 99.65 (PLS) | log<sub>e</sub> |
|  |  | CV | 68.30 | 1.92 (PLS) | 207.54 (ARN) | √ |

347 Transformation (Transf.): / = no transformation, log<sub>e</sub> = natural logarithm, √ = square-root

We classified the parameters presented in Tables 1 and 2 into two categories. The first category included 11 physical and biotic factors used as a proxy for the natural variability of environmental conditions (lagoon surface, lagoon-sea connection, station depth, β_MM_, OMC, mean and CV of Sal, Temp, O_2_ sat). The second category included 14 biogeochemical variables that either cause eutrophication (*i.e.* mean and CV of [TN], [NO_x_], [NH_4_ ^+^], [TP], [DIP]) or are the direct consequence of eutrophication (*i.e.* mean and CV of [TN], [TP], [Chla], turbidity) (Souchu et al., 2010). Because of technical problems, no [NO_x_] and [DIP] data were available for the stations BOL and PLS. Consequently, we imputed the missing mean and CV values for these two parameters at both stations with the ‘imputePCA’ function. This function estimates the missing parameters by first performing a principal component analysis on all the environmental variables and then by using the regularized fitted matrix. These steps are performed iteratively until convergence is reached (Josse and Husson, 2012).

#### Macrobenthic community structure

First, we calculated the mean macroinvertebrate abundance at the station scale by averaging over the three replicates and transformed it using a Hellinger transformation, to give more weight to common taxa and less weight to highly abundant, rare and under-sampled taxa (Legendre and Legendre, 2012). We then classified the 41 stations based on these transformed mean abundances, using K-means partitioning and the Calinski-Harabasz criterion with 100 iterations (‘cascadeKM’ function). Our goal was to highlight similarities between lagoons based on common species without focusing on rare taxa, as each coastal lagoon can present unique species assemblages (Basset et al., 2007), and to help with the result interpretation thanks to a simple classification. We tested for significant differences in the composition of our macrofauna groups using the analysis of similarity (ANOSIM) test with 9999 permutations and Euclidean distances (‘anosim’ function). Finally, we characterized each group using the indicator species metric (‘indval’ function, 9999 permutations). A high value of this metric for a given taxon associated to a given group means this taxon is consistently more present (fidelity) and abundant (specificity) in the stations composing that group than in stations composing the other groups (Dufrêne and Legendre, 1997). For this last analysis, we adjusted p-values using the False Discovery Rate correction method (FDR; Benjamini and Hochberg, 1995).

#### Taxonomy-based indices

To characterize the macrobenthic community structure, we calculated the commonly used M-AMBI (Multivariate AZTI Marine Biotic Index), the three indices composing the M-AMBI - species richness (SR), Shannon diversity index (H’) and AMBI (Muxika et al., 2007) - plus the inverse of Simpson’s index (N2) and the total macrofauna density. The AMBI is an index based on the classification of benthic species into ecological groups, from the most sensitive to disturbances (i.e. organic matter enrichment) to the most opportunistic ones (Borja et al., 2000). The raw (i.e. untransformed) macrofauna abundances were used to calculate all these indices. For AMBI and M-AMBI calculations, we removed high order taxa without an ecological group value (Anemonia and Polycladida), planktonic (Mysidae) and hard-substrate taxa (Serpula and Serpulidae) (Borja et al., 2005). We computed M-AMBI using the R function available in Sigovini et al. (2013) and we defined the good reference conditions as the highest values of SR and H’ (46 and 3.73, respectively) and lowest value of AMBI (0.31) measured in the dataset, irrespective of the WFD physico-chemical status described above. The results are perfectly similar whether the reference values used are those described above, those from the French government or those from the reference lagoons (see supplementary material 3). All indices were computed at the replicate level (calculation of M-AMBI at this scale does not affect the results, see supplementary material 4); the mean and standard deviation (SD) values were then calculated for each station. Finally, we tested for differences in total macrofauna density, SR, H’, N2, AMBI and M-AMBI between previously defined groups of stations (see section *Macrobenthic community structure*) using non-parametric Kruskal-Wallis rank sum tests (Breslow, 1970).

#### Linking macrobenthic communities and environmental variables

To assess how natural environmental variations correlated with anthropogenic pressures (explanatory variables) and how each explanatory variable correlated with macrobenthic indices, we built a correlogram showing the significant Spearman correlations (‘cor_pmat’ function) between all the raw variables. We also tested for differences in the variables considered (explanatory variables and macrobenthic indices) between lagoon-sea connection levels using non-parametric Kruskal-Wallis rank sum tests (Breslow, 1970), followed by pairwise Wilcoxon tests for multiple comparison, while adjusting p-values with FDR (Benjamini and Hochberg, 1995). Then, we transformed (natural logarithm or square root transformation), when necessary, the environmental variables and macrobenthic indices to approximate normal distributions and reduce data spread, before standardizing all the explanatory variables to facilitate the interpretation of the multiple linear regression coefficients (Schielzeth, 2010). We log_e_-transformed the lagoon surface, station depth, SR, N2 and macrofauna density (see Table 2 for the rest of the transformations). To limit multicollinearity and avoid artificially inflating the fit of linear models, we reduced the number of environmental variables (see section *Environmental variables*) for all future analyses by considering Pearson correlations < |0.6|, corresponding to a p-value < 5. 10^-5^ and leading to 19 remaining environmental variables.

To determine which environmental variables most strongly influence lagoon macrobenthic communities, we performed a redundancy analysis (RDA, ‘rda’ function) between the 19 previously selected environmental variables and the Hellinger-transformed mean taxa abundances. We computed the variance inflation factor (VIF, ‘vif.cca’ function) to check for multicollinearity and sequentially removed variables with a VIF > 5 (James et al., 2013). We verified the significance of the full RDA model using the ‘anova.cca’ function (1000 steps) and we performed a forward model selection procedure (‘ordiR2step’ function) based on the adjusted R2 of the full model and a p-value of 0.05 (9999 permutations) (Blanchet et al., 2008). We then computed the marginal effect of each selected explanatory variable as the adjusted R² of the constrained ordination containing only the given variable as the predictor. The conditional effect of a variable was equal, during the forward selection procedure, to the additional amount of variance in species communities explained by the corresponding variable at the time it was included into the model. Finally, we performed a partial RDA using the variables previously selected, to isolate the impact of eutrophication from natural environmental variability on lagoon macrobenthic communities (Legendre and Legendre, 2012).

In the same way as for the constrained multivariate analysis, we implemented a multiple linear regression approach (without any interactions between variables) to investigate which variables related to eutrophication and natural variability most strongly influenced the macrobenthic indices (SR, H’, N2, total macrofauna density, AMBI and M-AMBI). As there were still 19 potential explanatory variables, we performed a preliminary selection by including all the quantitative variables which alone had a significant effect on the index as first or second-degree orthogonal polynomial and the lagoon-sea connection if at least one of the three categories (weak, moderate, strong) had a significant effect on the index (‘lm’ function). We included second-degree polynomials to account for potential non-linear relations (Pérez-Ruzafa et al., 2007b) and verified that the selected variables had VIFs inferior to 5. We also checked the model hypotheses using the residuals vs fitted values plot for linearity, the Shapiro-Wilk test for residual normality, the Breush Pagan test for residual homoscedasticity (‘ols_test_breusch_pagan’ function) and the ‘acf’ function for the detection of residual spatial autocorrelation based on the station coordinates. Then, we ran a stepwise model selection using the ‘stepAIC’ function set in both directions and the BIC criterion. Finally, we checked the hypotheses of the selected model as previously explained, we investigated potential high leverage points (Cook’s distance > 0.5) and we reran the selected model without these points to verify its robustness. To compare the relative effect of each selected explanatory variable on a given macrobenthic index, we expressed the slope estimates of the standardized variables as effect sizes when the variable was quantitative and as mean standardized effects when the variable was categorical (Schielzeth, 2010).

Lastly, we implemented a variance partitioning procedure using the ‘varpart’ function, to assess how much community variation or macrobenthic index variation could be: (i) uniquely attributed to eutrophication when the effect of natural variability had been removed, (ii) uniquely attributed to natural variability when the effect of eutrophication had been removed or (iii) jointly attributed to both effects (Legendre and Legendre, 2012). Prior to this analysis, we reran the variable selection procedures implemented at the community and macrobenthic index levels considering alternatively only the eutrophication variables or only the variables related to natural variability. We then included the variables selected by the aforementioned method in the variance partitioning, which was based on partial RDAs or on partial linear regressions and we presented the results using the adjusted R² values.

## Results

### Inter-lagoon environmental variability

The dataset considered in this study encompasses a wide diversity of lagoons in terms of physico-chemical state according to the WFD and environmental parameters associated to natural variability and eutrophication (Tables 1 and 2). First, only lagoons with a strong (Leucate, Gruissan, Thau, Berre and Diana) or a moderate (La Palme and Bages-Sigean) connection to the sea were in a good physico-chemical state (green in Table 1). All the stations sampled in these lagoons presented mean [Chla] below 7 µg.L^-1^ and all of them except the ones located in the deep Berre and Diana lagoons (station depth > 5.8 m) presented a high beta diversity of macrophyte morphotypes (β_MM_ > 0.53). Five stations sampled in these lagoons presented the lowest value for a eutrophication related variable (DIA: mean [TN], BGS: mean [Chla] and mean Turb, LEN : mean [TP], LES: mean [DIP], THW: CV [DIP]). However, the THE station (Thau) had the most variable [NH_4_^+^] and the BRN and BRS stations (Berre) had the lowest mean O_2_ sat of the dataset. Seven lagoons were in a moderate (yellow in Table 1) or in a bad physico-chemical state (orange in Table 1) and were either moderately (Ponant, Vaine, Urbino) or weakly (Grand Bagnas, Vaccarès, Biguglia and Palo) connected to the sea. In this group, the GBA and VAC stations respectively had the lowest mean [NO_x_] and the lowest CV [TN] of the dataset whereas the URB and BIG stations respectively had the highest CV [NO_x_] and CV [Chla]. All the remaining weakly connected lagoons were in a poor physico-chemical state (Canet, Campignol, Vendres, La Marette, La Palissade, Bolmon). In this group, the CAN station had both the highest mean [TP] and [DIP] and the lowest CV [NH_4_^+^] and [NO_x_] of the dataset, while the CAM and BOL stations presented respectively the highest mean [NO_x_] and CV [Chla] of the dataset. Finally, all the lagoons between the cities of Sète and Montpellier (Figure 1) were in a poor condition, despite being strongly (Ingril North, Ingril South, Prévost) or moderately (Vic, Pierre-Blanche, Arnel, Méjean and Grec) connected to the sea. All the stations located in these lagoons except PBE, had OMC above 7% and up to 13%. In this group, the MEW station presented the highest mean [Chla] and [TN] of the dataset while the GRE station had the highest mean [NH_4_ ^+^], the ARN station the highest CV [TP] and Turb and the INN station the highest CV [DIP] of the dataset.

Based on the significant Spearman correlations between the environmental variables (Figure 2), stations in the larger lagoons were significantly deeper, cooler in summer and less variable in terms of summer salinity, presented higher β_MM_ and lower values for 8 eutrophication related variables (*e.g.* mean [Chla], mean [NO_x_] and CV [NO_x_]). The deepest stations were also the ones with the highest OMC. Two variables related to natural environmental variability were also significantly different according to the lagoon-sea connection level (KW test, *p* < 0.05 and post-hoc pairwise Wilcoxon tests *p* < 0.05): mean Sal (moderate > weak and strong > weak) and CV Sal (weak > moderate > strong).

**Figure 2.**
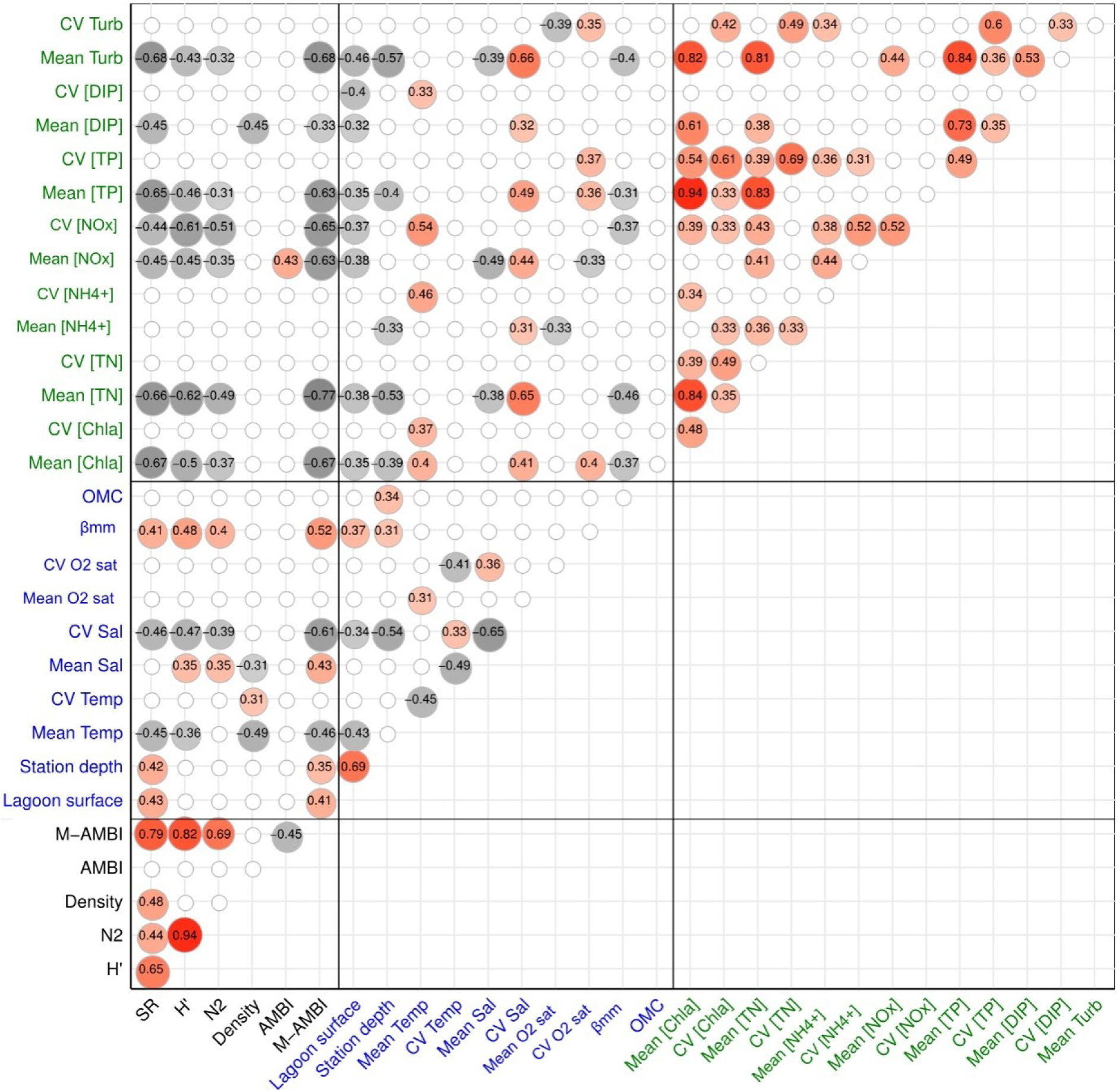
Correlogram showing only significant positive (red toned circles) and negative (grey toned circles) Spearman correlations (*p* < 0.05) between the untransformed taxonomic indices (in black, SR: species richness, H’: Shannon diversity, N2: inverse of Simpson’s dominance, M-AMBI: Multivariate AMBI), the untransformed environmental variables related to the lagoon’s natural variability (in blue, β_MM_: beta diversity of the macrophyte morphotypes, OMC: sediment organic matter content) and the untransformed environmental variables related to eutrophication expressed as means and coefficient of variations (in green, [Chla]: chlorophyll *a* concentration, [TN]: total nitrogen concentration, [NH_4_ ^+^]: ammonium concentration, [NO_x_]: nitrate plus nitrite concentration, [TP]: total phosphorous concentration, [DIP]: dissolved inorganic phosphorous concentration) calculated across 41 stations located in 29 lagoons along the French Mediterranean coast.

### Macrofauna communities and macrobenthic indices across French Mediterranean lagoons

A total of 42 465 benthic macroinvertebrates were counted and identified across all samples, belonging to 227 different taxa with most organisms identified to species level (74.5%), others to genus (17.2%), family (2.6%), order (2.6%), subclass (0.4%), class (1.8%) or phylum (0.9%). Across the 227 taxa, 1.3% were identified in over 50% of the stations, 90% in less than eight stations and 40% in only one station. No taxon was present in all 41 stations. Polychaetes were by far the most diversified group (102 taxa), followed by Malacostraca crustaceans (54 taxa), bivalves (25 taxa) and gastropods (19 taxa). Among the three taxa with a frequency of occurrence above 50%, two were polychaete species (*Hediste diversicolor* (66%) and *Heteromastus filiformis* (51%)), and the third was the bivalve *Cerastoderma glaucum* (51%). These three species also presented an important inter-station variability in abundance. For all the following analyses, we considered 217 taxa (see section *Benthic macroinvertebrates*), except for the AMBI and M-AMBI calculation, for which we considered only 213 taxa (see section *Taxonomy-based indices*).

The K-means clustering delineated two groups of stations with distinct macrobenthic communities (minimal difference in Calinsky-Harabasz criterion = 1.4), a result confirmed by the analysis of similarity test (ANOSIM statistic = 0.51, *p* < 0.001). These groups did not correspond to any obvious spatial or salinity pattern, as exemplified by the four Corsican stations and the seven oligo-mesohaline stations which were classified in both groups (Figure 1). The polychaete *Hediste diversicolor* (*p* = 0.022) was an indicator taxa of group A stations (red in Figure 1) along with the bivalve *Abra segmentum* (*p* = 0.022), Chironomidae (*p* = 0.029) and the amphipod *Gammarus aequicauda* (*p* = 0.038). Conversely, group B stations (blue in Figure 1) were significantly characterized by the bivalve *Scrobicularia cottardii* (*p* = 0.048), closely followed by another bivalve, *Loripes orbiculatus* (*p* = 0.065).

Finally, we found significant differences among the two groups in terms of SR and M-AMBI; group A was composed of stations with lower SR and weaker M-AMBI values than group B (SR: χ^2^(1) = 6.41, *p* = 0.011; M-AMBI: χ^2^(1) = 5.76, *p* = 0.016). Group B was also characterized by stations with higher H’ and N2 values and higher maximal macroinvertebrate densities than stations composing group A, albeit these tendencies were not significant (Appendix 3).

### Relationship between macrofauna communities and environmental variables

In all the following analyses, six variables were not considered because they were highly correlated with another variable (|r| ≥ 0.6 ; *p* < 5. 10^-5^): station depth (to lagoon surface), CV Sal (to mean Sal), mean [TN], mean [TP], mean Turb (to mean [Chla]) and CV [TP] (to CV [TN]). Redundancy analysis (RDA) was used to relate the macrofauna communities recorded within the lagoons to their natural environmental variability and their eutrophication levels (Figure 3). Mean Sal was also not considered because of a VIF > 5 linked to significant differences between lagoon-sea connection levels (see section *Inter-lagoon environmental variability*). The final RDA model, determined using a forward model selection procedure, explained 22% of the inter-station community variability (model adj. R^2^) and included six variables related to natural environmental variability (lagoon-sea connection level (*p* < 0.001), mean Temp (*p* < 0.001), lagoon surface (*p* = 0.0076), CV Temp (*p* = 0.013), mean O_2_ sat (*p* = 0.026) and OMC (*p* = 0.041) and two eutrophication related variables (mean [Chla] (*p* < 0.001) and mean [DIP] (*p =* 0.024)). Regarding the individual effect of environmental variables, mean [Chla] (7.22%), lagoon surface (5.90%), lagoon-sea connection (5.30%) and mean Temp (5.04%) were the most influential variables according to their marginal effect. Based on conditional effects, the most influential variables were mean [Chla] (7.22%), lagoon-sea connection (4.64%), mean Temp (3.62%) and lagoon surface (1.69%) (Appendix 4).

**Figure 3.**
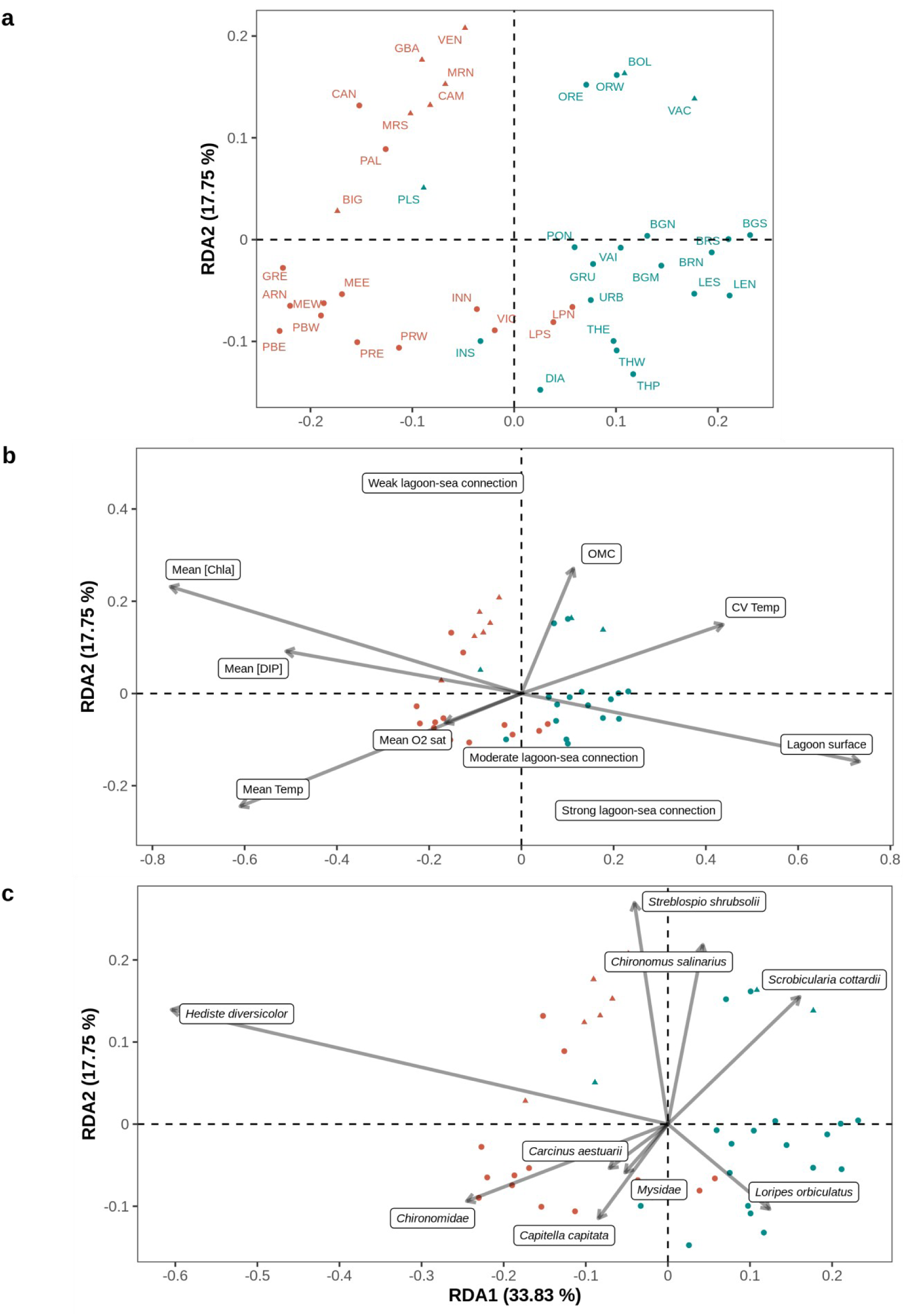
Redundancy analysis plots (RDA, axis 1 and 2) focusing on (a) the benthic macrofauna sampled across 41 stations located in 29 lagoons along the French Mediterranean coast (see Table 1 for the full lagoon names and corresponding stations) as explained by the set of environmental variables related to eutrophication and natural variability composing the most parsimonious model, (b) the selected environmental variables, where mean [Chla], lagoon surface, mean O_2_ sat and mean [DIP] were log_e_ transformed and CV Temp was square-root transformed (see Tables 1 and 2 for the full environmental variable names and values) and (c) the macrofauna taxa, where only taxa with a goodness of fit (cumulative proportion of variance explained by the first two axis) above 0.25 are plotted. In (a), inter-station distances are euclidean while in (b) and (c) angles between all vectors reflect linear correlations. Across all plots, each station (point) is represented using a symbol for its salinity type (triangle: oligo- and meso-haline and circle: poly- and eu-haline) and a color (red: group A and blue: group B) for its group membership (K-means partitioning) based on Hellinger-transformed mean abundances.

The first two axes accounted for 33.8 and 17.8% of the RDA model variance, respectively. First, the stations from group A and B were clearly separated along the first axis of the RDA, except for LPN, LPS, PLS and INS (Figure 3a). On the ordination of the environmental variables (Figure 3b), lagoon surface and CV Temp grouped on the positive side of the first axis while mean [Chla], mean [DIP] and mean Temp grouped on the negative side of the first axis. Consequently, group B macrobenthic communities were linked to large lagoons (also relatively deep), several of which presented high summer temperature variations. They were characterized by higher relative abundances of the bivalves *S. cottardii* and *L. orbiculatus* (Figure 3c). Conversely, group A communities were associated to smaller lagoons, subject to higher mean summer temperatures and high eutrophication levels. They were characterized by higher relative abundances of the polychaete *H. diversicolor* and Chironomidae (Figure 3c).

The second axis mainly corresponded to a confinement and also to an organic matter content gradient, mostly separating each group into two sub-groups with the lagoons weakly connected to the sea presenting sediments enriched in organic matter located towards the positive side of the second axis and the moderately and strongly connected lagoons located towards the negative side of the second axis. As mean Sal and CV Sal were correlated to lagoon-sea connection level (see section *Inter-lagoon environmental variability*), this axis also corresponded to a salinity gradient with low and variable salinity stations characterized by a higher abundance of *Chironomus salinarius* and *Streblospio shrubsolii* (Figure 3c). All the oligo- and meso-haline stations (triangles on Figure 3) grouped on the upper part of the RDA space and most poly- and eu-haline stations (circles on Figure 3) grouped on the lower part of the figure. Furthermore, communities of lagoons weakly connected to the sea were characterized by higher abundances of the bivalve *Scrobicularia cottardii* for group B, while communities from lagoons moderately to strongly connected to the sea were characterized either by higher abundances of *L. orbiculatus* for group B or by a diversified suite of species like the decapod *Carcinus aestuarii* or the polychaete *Capitella capitata* for group A.

Once the effect of natural environmental variability on macrofauna communities was removed through partial RDA and the only explanatory variables left to vary were mean [Chla] and mean [DIP] (eutrophication variables), no differences could be made between the communities of groups A and B (Appendix 5). This partial RDA explained only 6.4% of the inter-station community variability (model adj. R^2^).

### Relationship between macrobenthic indices and environmental conditions

Based on the significant Spearman correlations between the raw environmental variables and raw macrobenthic indices (Figure 2), SR, H’, N2 and M-AMBI were significantly higher at stations where mean Temp was lower, salinity was more stable and eutrophication related variables were lower, especially mean [Chla]. Furthermore, H’, N2 and M-AMBI were significantly higher at stations where mean Sal and β_MM_ were higher while SR and M-AMBI were also significantly higher at deeper stations located in larger lagoons. AMBI appeared to be significantly correlated positively only with mean [NO_x_] while total macrofauna density was significantly higher at stations with lower but more variable summer water temperatures, lower salinity and lower mean [DIP].

The results of the multiple linear regressions identifying which variables influenced the macrobenthic indices are presented in Table 3. The most parsimonious models contained between two and four predictors and explained between 27% and 74% of the macrobenthic indices’ variability. All the models contained at least one predictor associated to eutrophication and at least one predictor associated to natural variability except the N2 and AMBI models, which only contained predictors associated to natural variability and eutrophication respectively. Lagoon-sea connection was selected in three models, H’, N2 and M-AMBI, and in all three, the response variable was significantly higher in strongly connected lagoons. Differently, H’ and N2 were significantly higher and lower in moderately connected lagoons, respectively, while M-AMBI was significantly lower in weakly connected lagoons. Mean Temp and mean Sal were selected in two models, SR and macrofauna density. Mean Temp negatively influenced SR and macrofauna density while mean Sal positively influenced SR and had a quadratic effect (concave form) on macrofauna density. β_MM_ was selected in two models, H’ and N2, and positively influenced both response variables. The AMBI model was composed of mean [TN] (positive effect) and CV [NH_4_ ^+^] (quadratic effect, concave form). Finally, three other eutrophication related variables, CV [TN], mean [NH_4_^+^] and CV [NO_x_], were selected respectively in H’, macrofauna density and M-AMBI models where they had either a negative effect (M-AMBI) or a quadratic effect (concave form, H’ and density).

**Table 3.**
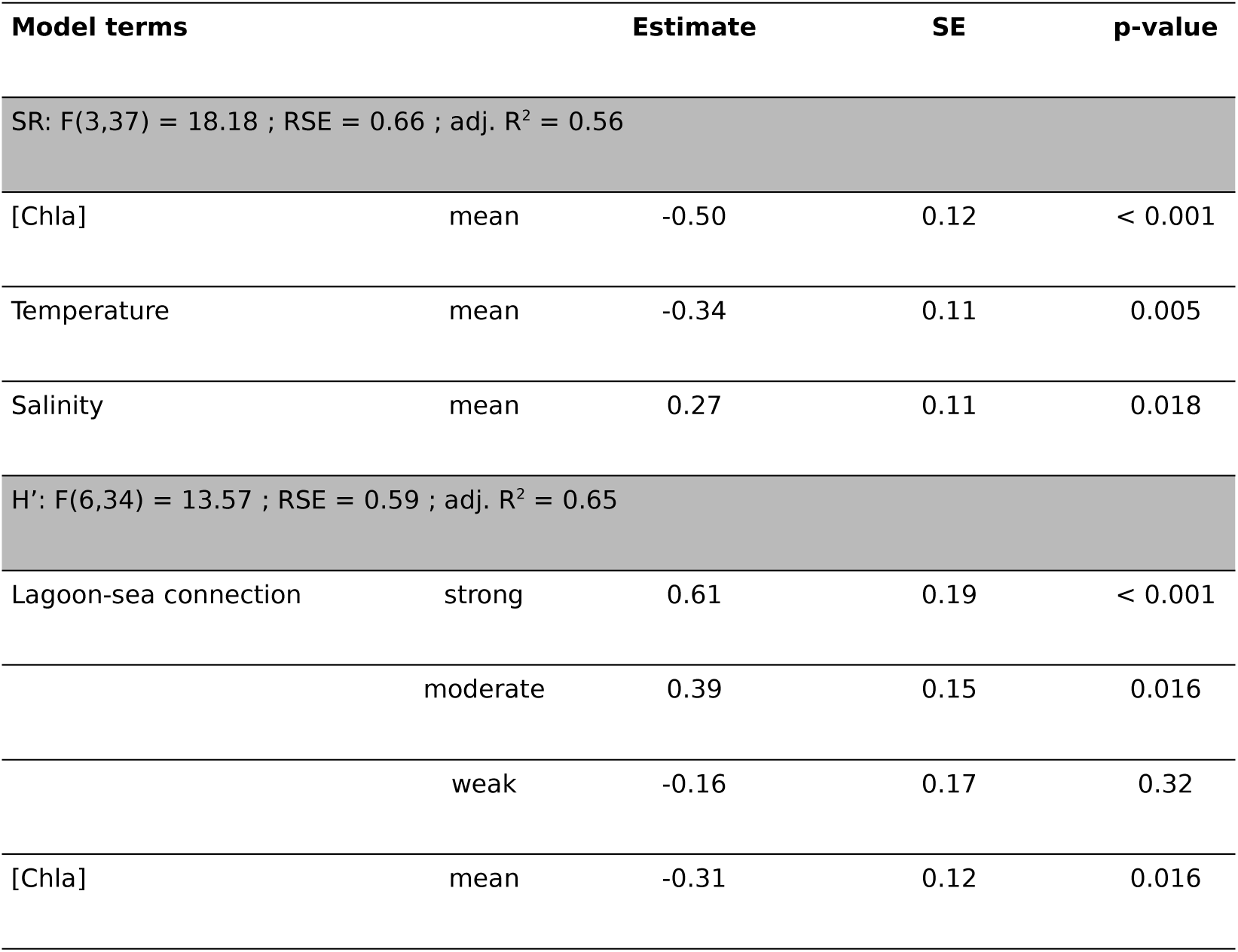
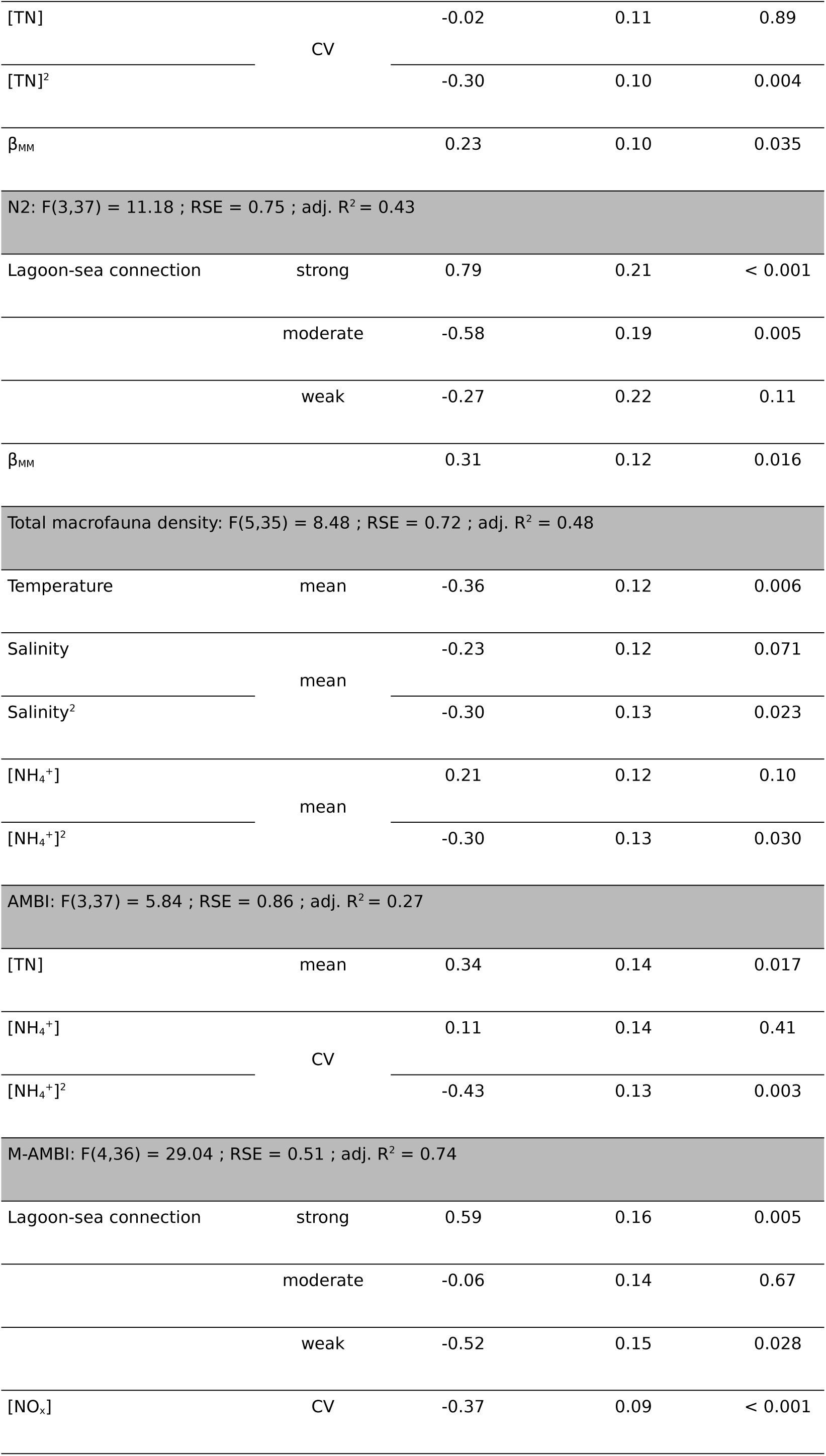
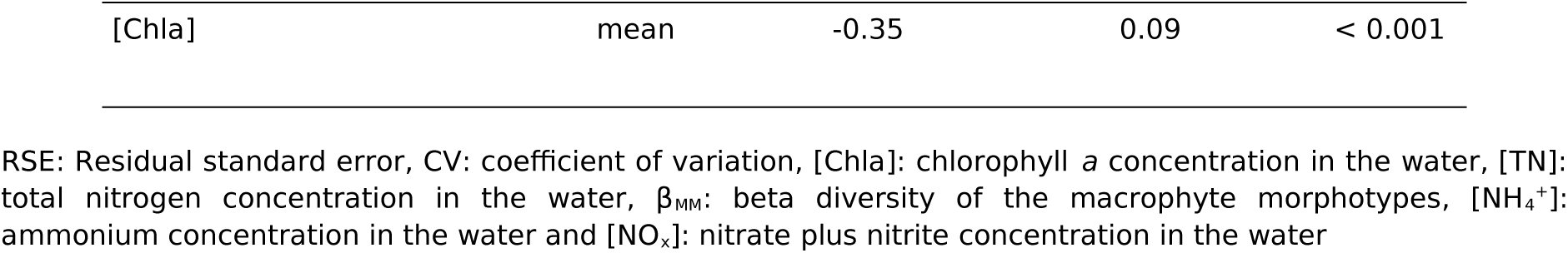
Most parsimonious models (ordinary least-square regressions) explaining the standardized macrobenthic taxonomic indices (species richness (SR), Shannon diversity (H’), inverse of Simpson’s dominance (N2), AZTI’s Marine Biotic Index (AMBI), Multivariate AMBI (M-AMBI)). SR, H’, N2 and total macrofauna density were log_e_ transformed prior to standardization. Both eutrophication related and natural variability related variables are considered as explanatory variables and the slope estimates are expressed either as effect sizes when the variable is quantitative or as mean standardized effects when the variable is categorical.

To isolate the independent effect of eutrophication on macrobenthic indices, we also plotted the partial effect of mean [Chla] on SR, H’ and M-AMBI once the effect of the other predictors retained in the minimal adequate models (Table 3) had been removed (Appendix 6). After controlling for the effect of mean Sal and mean Temp, the simple linear regression model between mean [Chla] and SR was still significant (adj. R^2^ = 0.32, F(1, 39) = 19.55, *p* < 0.001). It was also the case for the model between mean [Chla] and H’ after controlling for the effect of lagoon-sea connection, CV [TN], CV [TN]^2^ and β_MM_ (adj. R_2_ = 0.14, F(1, 39) = 7.33, *p* = 0.01) and for the model between mean [Chla] and M-AMBI after controlling for lagoon-sea connection and CV [NO_x_] (adj. R^2^ = 0.26, F(1, 39) = 15.41, *p* < 0.001).

To synthesize our results, we plotted for each station the M-AMBI along with the three macrobenthic indices composing it (SR, H’ and AMBI) (mean ± SD) as a function of mean [Chla] (Figure 4). Mean [Chla] was chosen as a synthetic indicator of eutrophication because it is (*i*) significantly and positively correlated to 9 out of the 13 eutrophication related variables (Figure 2), (*ii*) the eutrophication related variable (with mean [TN]) that presents the strongest correlations with SR, H’ and M-AMBI (Figure 2), (*iii*) the most frequently selected eutrophication related variable in the macrobenthic index models (Table 3) and (*iv*) a relevant indicator of eutrophication status according to Bec et al. (2011) and Cloern (2001).

**Figure 4.**
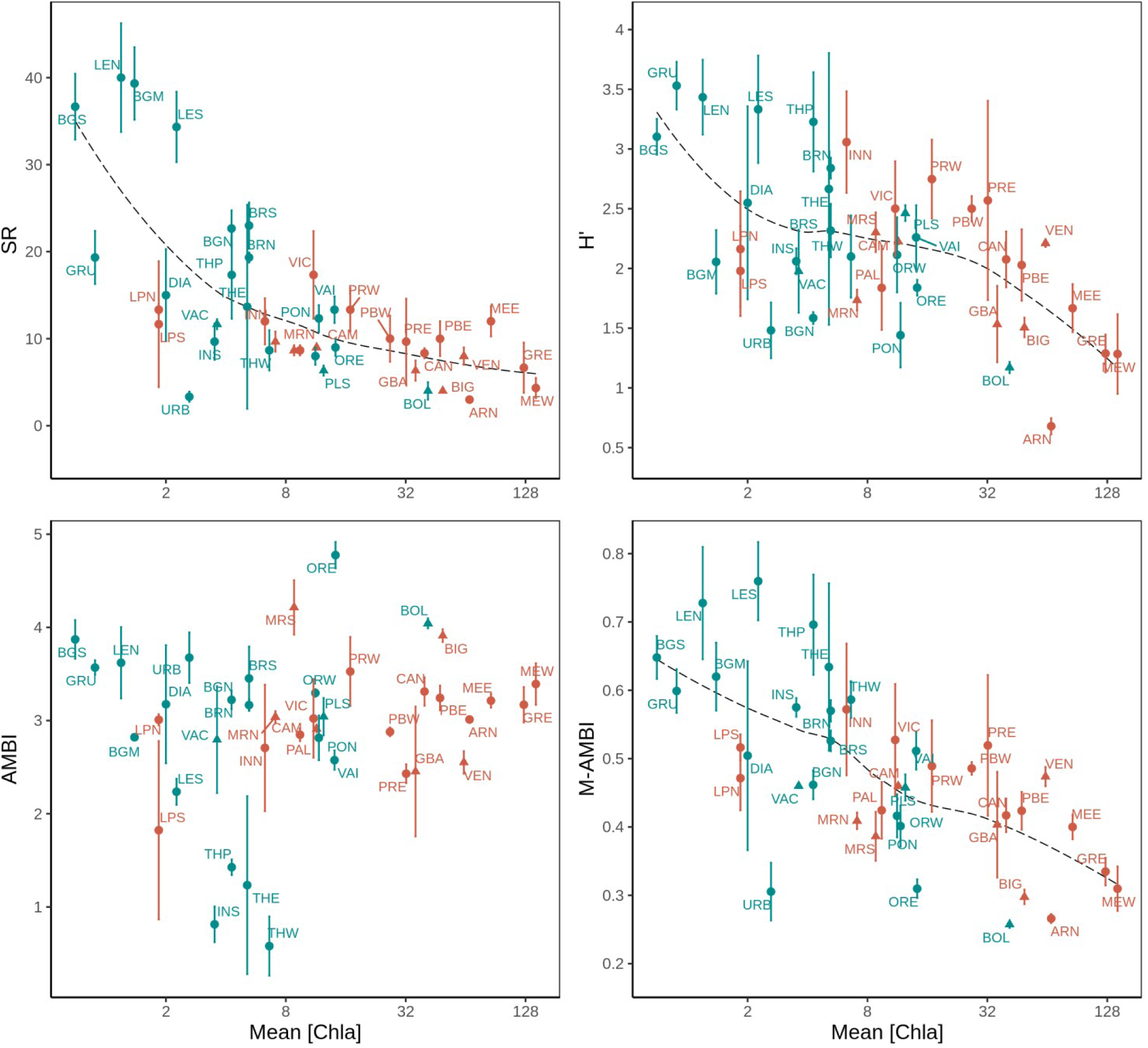
Mean (± SD) of the species richness (SR), Shannon diversity (H’), AMBI and M-AMBI indices as a function of mean chlorophyll a concentration (log_e_ scale) across 41 stations located in 29 lagoons along the French Mediterranean coast (see Table 1 for the full lagoon names and corresponding stations). Across all plots, each station (point) is represented using a symbol for its salinity type (triangle: oligo- and meso-haline and circle: poly- and eu-haline) and a color (red: group A and blue: group B) for its group membership (K-means partitioning) based on Hellinger-transformed mean abundances. The dashed lines correspond to loess regressions.

SR, H’ and M-AMBI were significantly correlated to mean [Chla] (r (Spearman) > | 0.50|) while AMBI was not (r = 0.12). Based only on the mean [Chla], most stations from group B (in blue) grouped towards higher levels of mean SR, H’ and M-AMBI than stations from group A (in red). Nonetheless, we identified an intermediary zone corresponding to mean [Chla] between 6.3 µg.L^-1^ and 14.2 µg.L^-1^ and composed of stations from both group located between the INN (SR = 12 ± 3, H’ = 3.06 ± 0.42 and M-AMBI = 0.57 ± 0.1) and ORE (SR = 9 ± 1, H’ = 1.84 ± 0.07 and M-AMBI = 0.31 ± 0.01) stations. In the line with these observations, the LPN, LPS and BOL stations appeared as outliers, as did LPN and LPS in the RDA (Figure 3). Finally, mean [Chla] was also significantly correlated (r (Spearman) = -0.40, *p* = 0.009) to the intra-station SR variability expressed using the standard deviation.

### Disentangling natural from anthropogenic factors

Based on the eutrophication and natural variability variables independently selected, we performed a variance partitioning, to investigate the amount of community and macrobenthic index variability uniquely linked to each variable category (Figure 5). Macrobenthic community and indices can be grouped into three categories according to the relative importance of natural variability alone, eutrophication alone, and the two together, in explaining their respective variability. (*i*) Macrofauna density was first influenced uniquely by natural variability (29.0%), then jointly by natural variability and eutrophication (11.5%) and finally uniquely by eutrophication (7.9%). (*ii*) Macrobenthic community, N2 and H’ were first jointly influenced by natural variability and eutrophication (community: 8.2%, H’: 33.6%, N2: 22.1%), and then uniquely by natural variability (community: 8.0%, H’: 17.8%, N2: 21.1%) and finally uniquely by eutrophication (community: 3.7%, H’: 13.9%, N2: 2.9%). (*iii*) SR and M-AMBI were first jointly influenced by natural variability and eutrophication (SR: 30.7%, M-AMBI: 49.0%), and then uniquely by eutrophication (SR: 20.3%, M-AMBI: 17.6%) and finally uniquely by natural variability (SR: 5.8%, M-AMBI: 8.9%).

**Figure 5.**
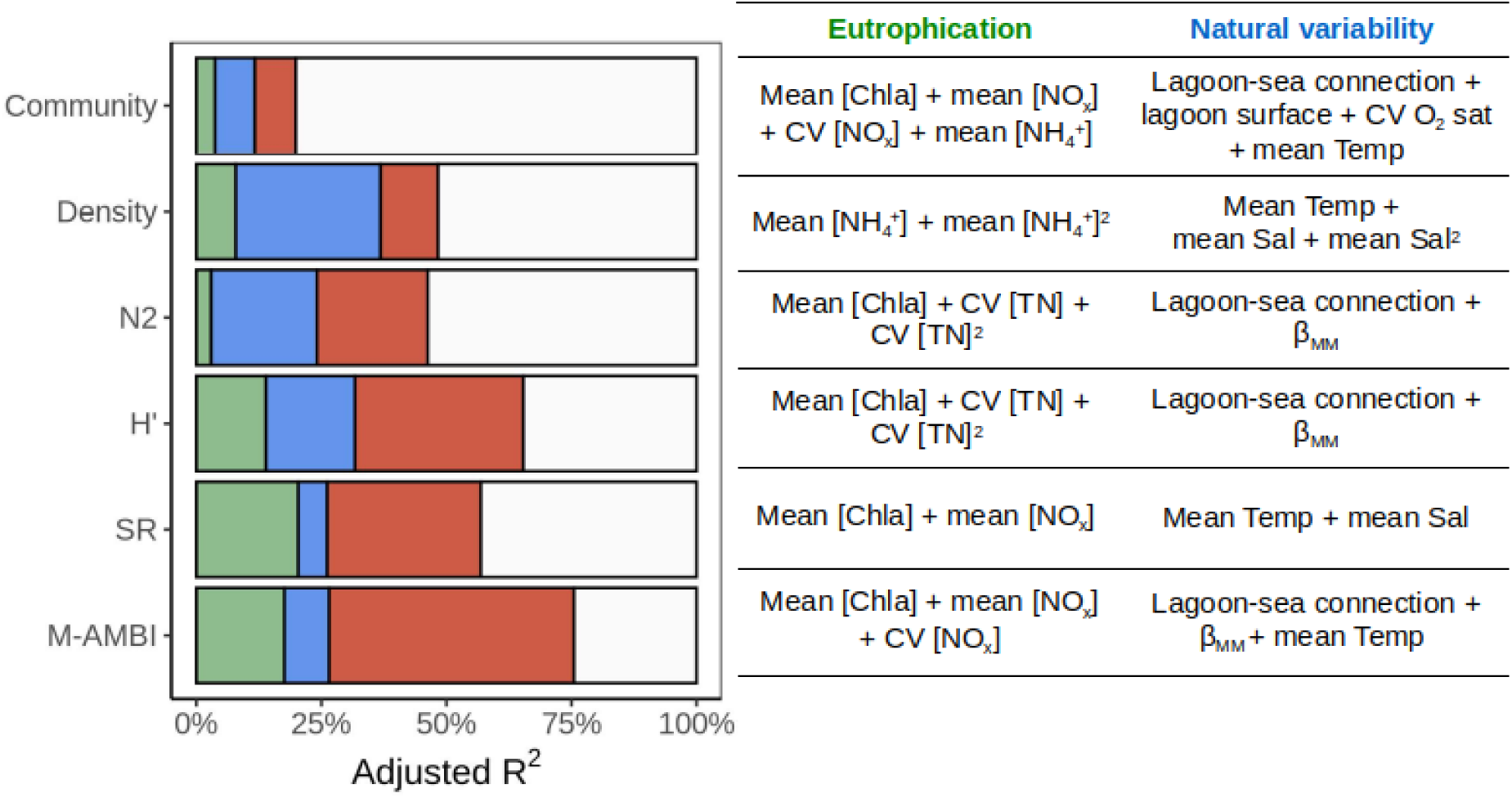
On the left, variance (adjusted R2) of the macrofauna community data (community), total macrofauna density (density), inverse of Simpson’s dominance (N2), Shannon diversity (H’), species richness (SR) and multivariate AMBI (M-AMBI) uniquely attributed to eutrophication (green), uniquely attributed to natural variability (blue) and jointly attributed to both effects (red). The residual variance is shown in white. On the right, eutrophication variables and variables related to natural variability considered in the variance partitioning of the macrofauna community and macrobenthic taxonomic indices.

Using figure 2, we can identify variables related to eutrophication or to natural variability that significantly co-vary and are jointly implicated in a macrobenthic index model (Table 3) and/or in a variance partitioning (Figure 5). Mean [Chla] was significantly correlated with lagoon surface (-0.35), β_MM_ (-0.37), mean Temp (0.40) and CV O_2_ sat (0.40). Mean [NO_x_] was significantly correlated with lagoon surface (- 0.38), mean Sal (-0.49) and CV O_2_ sat (-0.33) while CV [NO_x_] was significantly correlated with lagoon surface (-0.37), β_MM_ (-0.36) and mean Temp (0.53). Finally, mean [DIP] and lagoon surface were significantly correlated (-0.32). Lagoon-sea connection, a qualitative variable, was another source of confounding effects between natural variability and eutrophication on the macrobenthic indices and community. Indeed, there was a significant difference between the lagoon-sea connection and: *(i)* mean [Chla] (weak > strong connection); *(ii)* mean [NO_x_] (weak > strong connection) and (*iii*) CV [NO_x_] (moderate > strong).

## Discussion

As outlined by Cloern (2001), eutrophication is a complex phenomenon with direct (*e.g.* changes in dissolved nutrient contents) and indirect (*e.g.* changes in community) responses that are modulated by attributes of each recipient system. This study is a first step towards better understanding how hydro-morphology, natural environmental variability and eutrophication jointly shape macrobenthic communities in Mediterranean coastal lagoon ecosystems.

### Macrobenthic communities are driven by lagoon hydro-morphology and natural environmental (in)stability

Lagoon macrobenthic communities appear to be first determined by lagoon hydro-morphology and natural environmental stability. When investigating the independent effect of each environmental parameter on macrobenthic communities, lagoon-sea connection level was the variable explaining the most of H’, N2 and M-AMBI variability and the second most important variable explaining community composition. This qualitative variable is a proxy for marine water renewal (*i.e.* residence time), a gradient well known for characterizing coastal lagoon biological communities (Barbone and Basset, 2010; Perez-Ruzafa and Marcos-Diego, 1992; Tagliapietra et al., 2009), and closely linked to mean Sal (moderate > weak and strong > weak) and CV Sal (weak > moderate > strong). Salinity, historically used to classify coastal lagoons (Anonymous, 1959), was also an important driver of SR (linear positive effect) and total macrofauna density (concave effect). Indeed, simple and multimetric descriptors of lagoon macrobenthic communities like species richness are known to be highly and positively influenced by mean water salinity (Barbone et al., 2012; Magni et al., 2022; Pitacco et al., 2019; Reizopoulou et al., 2014). Differently from us, Barbone et al. (2012) reported a macrofauna density decrease from oligo- to eu-haline lagoons, maybe because only undisturbed lagoons were considered in their study. Furthermore, the effect of lagoon-sea connection on N2, a diversity index sensitive to high abundance taxa (Morris et al., 2014), indicates that a strong connection to the sea promotes benthic communities with significantly more relatively abundant taxa whereas moderate and weak connections promote communities dominated by a few abundant and in our case, euryhaline taxa (*e.g. Hediste diversicolor, Streblospio shrubsolii*), a characteristic typical of naturally stressed environments like lagoons subject to strong salinity variations (Elliott and Quintino, 2007). This result seems to confirm that moderately and weakly connected lagoons are highly selective environments for benthic organisms at least partly because of their strongly fluctuating environmental conditions (Barnes, 1980; Provost et al., 2012; Zaldívar et al., 2008). The negative effect of decreasing salinity and lower connection levels to the sea on SR, N2 and macrofauna density can also be linked to the physiological stress that natural and abrupt salinity variations (*e.g.* droughts, heavy rainfall) can represent for larvae, juveniles and adults of marine-originating benthic organisms inhabiting coastal lagoons (Cognetti and Maltagliati, 2000; Magalhães et al., 2019; Vernberg, 1982). Overall and based on other published evidence, our results could be explained by (*i*) water renewal from marine origin, (*ii*) primary colonization and/or post-disturbance recolonization of lagoons by marine-originating larvae through dispersal and recruitment and (*iii*) environmental (in)stability, which are known to strongly shape lagoon benthic communities (Basset et al., 2006; Pérez-Ruzafa et al., 2007b; Pérez-Ruzafa and Marcos-Diego, 1992), as proposed in the confinement theory (Guelorget and Perthuisot, 1983).

Water temperature was another important driver of lagoon macrobenthic communities, SR and macrofauna density in our study. Temperature had a negative effect on these communities, probably directly through thermal stress, and indirectly through its effect on oxygen dynamics. Higher and more variable summer temperatures directly represent an abiotic constraint for benthic organisms by increasing their oxygen demand and reducing their mobility (Vernberg, 1982). At the same time, high summer temperatures decrease oxygen solubility and increase the probability of hypoxia and anoxia (Derolez et al., 2020a) while also reducing the tolerance of benthic organisms to low dissolved oxygen (Vaquer-Sunyer and Duarte, 2011). This cocktail of effects can ultimately cause the death of sensitive taxa, such as large sized and long-lived species (Quillien et al., 2015; Reizopoulou and Nicolaidou, 2007; Sigala et al., 2012) and reduce the ability to move of more tolerant taxa (Vernberg, 1982), hence increasing their susceptibility to predation and decreasing their possibility to search for food and/or for less stressful environmental conditions (Díaz and Rosenberg, 1995; Rosenberg et al., 1992; Weissberger et al., 2009).

The diversity of macrophyte morphotypes (β_MM_) also affected positively H’ and N2, and was positively correlated to all macrobenthic indices except density and AMBI. Simple taxonomic indices like lagoon macrobenthic species richness, biomass density and numerical density are locally sensitive to vegetation presence and type (Arocena, 2007; Barbone et al., 2012, Magni and Gravina, 2023). Conversely, we show here the probable effect of benthic habitat heterogeneity expressed at the lagoon or sub-lagoon scale on the structure of benthic communities at a local scale (*ca.* 50 m^2^ station), an effect mainly visible when focusing on intermediate and high abundance taxa (Morris et al., 2014). Focusing on macrobenthic fauna at the lagoon scale, β_MM_ informs us on the number of potential (*i*) environmental niches that can be filled by the species pool and (*ii*) refuges (Magni and Gravina, 2023; Nordström and Booth, 2007; Ware et al., 2019) from which these organisms can (re)colonize habitat patches after mortality events caused by temporary unfavorable conditions. A model of local extinctions linked to adverse environmental conditions, followed by recolonization events via pelagic larvae was suggested by Vergara-Chen et al. (2013) to explain the small-scale genetic structure of the cockle *Cerastoderma glaucum* inside the Mar Menor lagoon (Spain). The positive effect of β_MM_ on macrobenthic diversity also provides us with a few leads on how macrophytes can structure lagoon macrobenthic communities. Macrophytes can act as (*i*) temporary refuges from vagile predators for small mobile benthic organisms like amphipods (Nordström and Booth, 2007; Ware et al., 2019), (*ii*) a food source for grazers and detritus-feeders (Carlier et al., 2007; Lepoint et al., 2000; Ouisse et al., 2011; Vizzini and Mazzola, 2006) and (*iii*) settlement promoters for pelagic particles like phytoplankton and larvae (Colden et al., 2016; Donadi et al., 2014; González-Ortiz et al., 2014), especially in the case of meadow-forming species like *Zostera.* All these local effects are likely to spill-over to adjacent unvegetated sediments (Boström et al., 2006; Magni et al., 2017).

Overall, natural environmental (in)stability, connection level with adjacent water systems and/or number of potential environmental niches seem to be the main drivers of lagoon macrobenthic communities and to be stronger drivers of benthic macrofauna assemblages than eutrophication, which probably represents an additional source of stress for benthic invertebrates (Barbone and Basset, 2010).

The environmental variables considered in the manuscript do not unfortunately explain the communities described at the LPN and LPS stations. These two macrofauna stations sampled were actually classified in a group generally associated to small eutrophic lagoons and mainly composed of stations sampled in lagoons evaluated by the WFD as being in a poor or in a bad physico-chemical status even through is considered in a good physico-chemical status according to Chla concentration. This could be the result of the strong variations in salinity recorded throughout the year, which we have not been considered here. In this lagoon, marked annual salinity variations are observed (between 3.6 and 76.8 in Wilke and Boutiere (2000)), variations indetectable with the available summer salinity data. Consequently, the eight environmental variables selected to explain the overall macrofauna communities were not able to explain the one characterizing this lagoon. This result also reinforces the importance of natural environmental instability probably has on structuring benthic macrofauna communities.

### Eutrophication affects lagoon macrobenthic communities through interconnections with natural environmental variability

Eutrophication appears to act as an additional disturbance for lagoon macrobenthic communities via direct (*e.g.* toxicity of inorganic nitrogenous compounds) and indirect mechanisms (*e.g.* hypoxic stress). The three groups derived from the variance partitioning approach highlight that the most influential category of variables (*i.e.* natural variability alone, eutrophication alone, natural variability and eutrophication together) is not the same depending on the level of biological organization we focus on. Indeed, the benthic invertebrate density was first explained by natural variability alone and more precisely by physiologically constraining variables (salinity and temperature) whereas macrobenthic community composition, SR, H’ and N2 were first explained jointly by natural variability and eutrophication. With these joint effects, we probably measure *(i)* how lagoon-sea connection modulates the susceptibility (*i.e.* propensity) of each lagoon to eutrophication, *(ii)* how higher water temperatures promote phytoplankton blooms and *(iii)* how eutrophication reduces benthic habitat heterogeneity, followed by their respective effects on benthic community diversity.

*(i)* Lagoon-sea connection informs us on each lagoon’s water residence time and consequently, on its retention potential regarding nutrients and pollutants (Chapelle et al., 2001; Cloern, 2001; Fiandrino et al., 2017). As such, watershed related nutrients and their direct effect on primary producers are buffered by each lagoon’s degree of connection to the sea, as indicated by the significantly higher mean [Chla] and mean [NO_x_] in lagoons weakly connected to the sea compared with strongly connected ones. *(ii)* Independently from lagoon hydro-morphology, the positive correlation between mean [Chla] and mean temperature illustrates how higher water temperatures increase phytoplankton growth rates which can lead to blooms (Rose and Caron, 2007), as recently demonstrated in Thau (Trombetta et al., 2019), a large lagoon strongly connected to the sea. Nonetheless, it seems that eutrophication is buffered in larger lagoons where water temperatures increase less in summer, as suggested by the negative correlation between lagoon surface and mean summer temperature. *(iii)* Furthermore, one of the direct responses of Mediterranean coastal lagoons to changes in nutrient inputs is an increase in pelagic primary production (Souchu et al., 2010), which in turn decreases water transparency (Cloern, 2001) and limits light availability for benthic primary producers (Le Fur et al., 2019, 2018). Ultimately, sustained eutrophication of coastal lagoons can cause a regime shift from a primary producer community dominated by perennial macrophytes presenting different morphotypes, to a community dominated by free-floating and opportunistic macroalgae like *Ulva rigida* and finally, to a phytoplankton-dominated community (Le Fur et al., 2019). The negative correlation between several eutrophication variables (*e.g.* mean [Chla], mean turbidity) and β_MM_ highlights the simplification of benthic habitats caused by lagoon eutrophication (Cloern, 2001; Le Fur et al., 2019, 2018) that shift from a mosaic of different macrophyte morphotypes to a sediment void of macrophytes or with one macrophyte morphotype. Overall, the multiple interconnections between natural environmental variability and eutrophication likely explain the high amount of macrobenthic community and index variance jointly attributed to both effects, as underlined in the conceptual model developed by Cloern (2001).

The unique effect of eutrophication variables on each taxonomic component also informs us about which of these variables likely have a direct effect on macrobenthic invertebrates without being modulated by natural variability. Indeed, benthic organisms can be affected by eutrophication directly through the toxicity of inorganic nitrogenous compounds like ammonium, nitrites and nitrates (Jessen et al., 2015). Our results indicate that high NH_4_^+^ concentrations seem to have a very small negative effect on total macrofauna density while NO_x_ toxicity could explain the strong unique effect of eutrophication on SR and M-AMBI, and both NH_4_^+^ and NO_x_ toxicity could be responsible for the very weak unique effect of eutrophication on macrobenthic communities. These results are probably also linked to indirect effects high eutrophication levels can have on macrobenthic communities via increased water turbidity and sediment bio-geochemistry modifications (*e.g.* anoxic superficial sediments (Zilius et al., 2015)). Indeed, water NH_4_^+^ and NO_x_ concentrations in excess can be the sign of highly eutrophic lagoons (Souchu et al., 2010). Overall, experimental work on a few well-selected species and populations sampled in lagoons presenting different levels of natural variability would greatly improve our understanding of the effect of eutrophication on lagoon benthic invertebrates via inorganic nitrogenous compound toxicity. Even trough marine animals appear to be less sensitive to nitrate than freshwater animals, nitrate toxicity to benthic invertebrate increases with increasing nitrate concentrations and exposure times (Alonso and Camargo, 2006; Boardman et al., 2004; Camargo et al., 2005) Nitrate toxicity may also increase with decreasing body size, water salinity, and environmental adaptation. In lagoon environments, where organisms are often smaller in size, this could also be of importance.

Furthermore, mean [Chla] appears to have a strong negative effect on macrobenthic communities by impacting the diversity of low and intermediate abundance taxa (i.e. SR and H’ are diversity indices that give more weight respectively, to low abundance/rare and intermediate abundance taxa) and this effect is at least partly independent from the selected natural variability variables (*i.e.* mean temperature, mean salinity for SR and lagoon-sea connection, β_MM_ for H’). Two non-exclusive hypotheses could explain this result: small-scale habitat homogenization and hypoxic stress. Despite not being the appropriate index to investigate beta diversity, the standard deviation of SR at the intra-station scale was significantly correlated negatively to mean [Chla], which seems to point towards a homogenization of benthic communities at the meter scale (i.e. a few meters separate each replicate of a given station) linked to eutrophication. The loss of benthic taxa presenting restricted environmental niches and dependent upon small-scale habitat diversity for reproduction, recruitment, refuge and/or food (Brauns et al., 2007; Harman, 1972; Stendera and Johnson, 2008) could explain this result. Secondly, low to zero dissolved oxygen (*i.e.* hypoxia and anoxia) is another well-known symptom of eutrophication in transitional waters like coastal lagoons (Chapelle et al., 2001; Cladas et al., 2016; Vignes et al., 2010). Oxygen depletion generally takes place during the summer and is maximal at the end of the night/early morning (Newton et al., 2010), often starting near the bottom because of benthic organic matter mineralization (*e.g.* sedimented phytoplankton, dead macroalgae) and low water renewal (Rigaud et al., 2021; Souchu et al., 1998). In the worst cases, these low to zero oxygen zones can spread to the entire water column and last for several days (Chapelle et al., 2001; Cladas et al., 2016), as seen during the 2018 dystrophic crisis in Thau lagoon (Derolez et al., 2020a). These crises are known to impact macrobenthic community first in terms of abundance and then in terms of composition and species richness depending on the sensitivity of each species to reduced oxygen (Díaz and Rosenberg, 1995; Riedel et al., 2012). Consequently, lagoon macrobenthic invertebrates are often reported as being less abundant and less diverse in fall than in spring because of summer hypoxic stress (Barbone et al., 2012; Basset et al., 2013a; Orro and Cabana, 2021). The negative impact of eutrophication via hypoxic/anoxic stress on lagoon macrobenthic communities is suggested by the RDA results where high mean O_2_ sat (*i.e.* hyper-saturation linked to strong pelagic primary production) is associated to group A stations that present significantly lower species richness than group B stations. Nonetheless, oxygen related variables were not selected in the other models, a result probably linked to when oxygen was measured (late morning) and at what depth it was measured (generally sub-surface). Indeed, these features limit our ability to detect potential short and long-term effects of hypoxic stress on macrobenthic communities. Bottom hypoxia/anoxia is also known to predominantly take place in deep lagoons like Berre and Thau (Derolez et al., 2020a; Rigaud et al., 2021; Souchu et al., 1998), where wind derived water mixing is limited by depth. Nonetheless, anoxic crises inside coastal lagoons are expected to increase in frequency and intensity with climate change (Derolez et al., 2020a; Miyamoto et al., 2019). Consequently, it appears paramount to (*i*) study the sensitivity of lagoon benthic organisms to low dissolved oxygen, not only of freshwater and marine taxa (Díaz and Rosenberg, 1995; Landman et al., 2005) and (*ii*) monitor more closely summer bottom dissolved oxygen concentrations, especially at higher frequency (O_2_ being very variable) and at temporal scales relevant to these organisms (*i.e.* across several days starting when bottom hypoxia is detected (Riedel et al., 2012)).

Overall, lagoon hydro-morphology (connection to the sea, surface and depth) which affects the level of environmental variability (salinity and temperature), as well as lagoon-scale benthic habitat diversity (β_MM_) seem to regulate the distribution of macrobenthic species (Bachelet et al., 2000), while eutrophication and associated stressors like low dissolved oxygen act upon the existing communities, mainly by reducing taxa richness and diversity (SR and H’). Disentangling the effects of anthropogenic stressors and natural environmental variables remains nevertheless challenging, especially as stressors and variables can co-vary, even in the absence of anthropogenic impact. Moreover, it is often impossible to measure effects of every single possible stressor on macrobenthic communites. In our study, grain-size could not be included to differences in laboratory protocols between lagoons and between years and heavy metal contaminations were not available at the same temporal and spatial scale (one measurement per lagoon every ten years). It should be relevant to re-run the analyses when such data would be available at the macrofauna sampling sites to test the effect of this supplementary anthropogenic driver to check if it would be discriminating benthic communities.

### The M-AMBI and recommendations for its use as an indicator for Mediterranean coastal lagoons

With this study, we investigate for the first time the joint sensitivity of the M-AMBI index to natural and anthropogenic stressors across 29 lagoons covering most of the French Mediterranean coastline. The M-AMBI (Muxika et al., 2007) is currently used to evaluate the WFD ecological state of poly- and eu-haline French Mediterranean lagoons based on soft sediment benthic invertebrate, with reference conditions using minimally impacted sites (European Commission, 2018). This index has previously been shown to be sensitive to pressures related to eutrophication, like concentration in chlorophyll *a* and total nitrogen (Derolez et al., 2014) but it is also known to be sensitive to natural environmental variability (*e.g.* water salinity) and lagoon hydro-morphology (*e.g.* surface and depth) (Barbone et al., 2012).

Our results show that the M-AMBI appears to be a sensitive indicator of eutrophication through the species richness and Shannon diversity components of the index but not the AMBI one. First, the strong correlation between M-AMBI and mean [Chla] was linked only to the significant correlation between SR and mean [Chla] and between H’ and mean [Chla] while AMBI was not significantly linked to most eutrophication related variables (except mean [NO_x_]) nor to chlorophyll *a* concentration, a common eutrophication indicator (Bec et al., 2011; Cloern, 2001). Importantly, these linear relations remained significant once the effect of identified covariables was removed. Regarding AMBI, its current application in transitional waters like coastal lagoons suffers from methodological limitations. Indeed, this index is based on macroinvertebrate sensitivity/tolerance to organic matter enrichment following Pearson and Rosenberg (1978) and was developed for coastal waters (Borja et al., 2000), not for coastal lagoons, which are known to sometimes present sediments naturally enriched in organic matter linked to the decay of meadow-forming macrophytes (Bachelet et al., 2000), watershed inputs and the overall low organic matter exportation (except during floods) (Perthuisot and Guelorget, 1983). These results highlight the limits of indirectly including AMBI in the evaluation of the ecological state of Mediterranean coastal lagoons. Nonetheless, its absence of sensitivity to natural variability and lagoon hydro-morphology suggests an index based on redefined ecological groups could be promising (Robertson et al., 2016), especially if we aim towards a more specific macrofauna-based index to evaluate eutrophication. Overall, increasing the robustness of M-AMBI would likely necessitate additional work to classify lagoon macroinvertebrates according to other eutrophication related stressors like low oxygen, nitrogenous compound toxicity and/or hydrogen sulfide toxicity.

Secondly, M-AMBI variability was two times more strongly attributed uniquely to eutrophication related variables than uniquely to natural variability ones, with nonetheless a very strong joint effect of the two types of variables. Our study reinforce the difficulty of M-AMBI to clearly identify natural disturbance but also the high sensitivity of this multimetric index to anthropogenic pressures such as eutrophication or metal contaminations and pesticides in transitional waters (Paul et al., 2023; Pelletier and Charpentier, 2023; Pollice et al., 2015). The strong joint effect is probably related, as previously discussed, to the same interconnections between lagoon-sea connection that buffers eutrophication, temperature that promotes phytoplankton blooms and eutrophication that homogenizes benthic habitats. Several modifications could help limit the sensitivity of M-AMBI to natural variability. First, as recommended by Borja et al. (2012) who pointed that the inability of this index to detect anthropogenic responses is often linked to the use of inappropriate reference conditions, the currently used poly- and eu-haline vs oligo- and meso-haline categories (Provost et al., 2012; European Commission, 2018) could be replaced by one based on the three lagoon-sea connection levels that strongly influence the M-AMBI or on another typology like the choked vs restricted typology (Barbone et al., 2012; Basset et al., 2013b), with the test of new reference lagoons (European Commission, 2018). Secondly, the replacement of a sampling done with an Ekman-Birge grab by a diver-operated sampling would limit heterogeneity between replicats by (i) verifying that only unvegetated sediments are being collected and (ii) describing the presence and type of macrophyte near the sampling site (*e.g.* 10 meter radius). This could represent a bias in many homogeneous environments where random selection of the sampling zone is quite possible but macrophytes are often heterogeneously distributed in coastal lagoon and areas of unvegetated sediments are rare. Selection and description of the sampling area by divers could therefore be relevant in these shallow environments even if this often means additional human and financial costs. This last modification would likely help reduce the effect of local benthic habitat heterogeneity on H’ and M-AMBI (Magni and Gravina, 2023; de Paz et al., 2008).

Finally, several studies have underlined the limits of using the M-AMBI based on abundance to evaluate the ecological state of transitional ecosystems like coastal lagoons, as opportunistic and tolerant species are generally numerically dominant in these systems (Mistri et al., 2018; Pitacco et al., 2019; Puente and Diaz, 2008). Other indicators based on the size spectra and biomass of benthic macroinvertebrates like the biomass based M-AMBI (M-bAMBI (Mistri et al., 2018)) and the index of size distribution (ISD (Reizopoulou and Nicolaidou, 2007)) have showed promising results and applying these indices to French coastal lagoons would represent an additional trial in the context of historically highly eutrophic lagoons currently undergoing a re-oligotrophication trajectory (Derolez et al., 2020a, 2020b; Le Fur et al., 2019; Leruste et al., 2016). Another interesting avenue would be to investigate more closely the spatial scales at which changes in macroinvertebrate abundance and diversity can be detected and link these spatial changes to localized anthropogenic disturbances like heavy metals (Magalhães et al., 2019) while also considering small-scale biotic interactions such as competition, predation and facilitation. Indeed, sediment contaminants like biotic interactions are likely to affect benthic macrofauna at fine spatial scales (*ca.* 1 km) that are more relevant to low mobility macroinvertebrates (Berthelsen et al., 2018; Brauko et al., 2020, 2015; Carvalho et al., 2006).

In a nutshell, to improve the robustness of M-AMBI, we call for a revision of the ecological groups at the base of the AMBI index and of the sampling protocol (*e.g.* using divers) along with a re-evaluation of the lagoon typology. More generally, it appears key to increase our understanding of the effects of other disturbances like hypoxia and heavy metals on lagoon invertebrates and to reevaluate the small-scale distribution of benthic organisms in French coastal lagoons, more than 30 years after the historical studies by Amanieu et al. (1977), Guelorget et al. (1994) and Guelorget and Michel (1979).

### Overcoming taxonomic limitations

A large part of the identified macroinvertebrates had a very restricted geographical range, preventing us from building a general species-based framework linking eutrophication and macrobenthic communities. Indeed, we found a very high level of lagoon-specificity for benthic macroinvertebrates across French Mediterranean coastal lagoons with no taxon present across all 41 stations, 40% of the 227 taxa identified in only one station and only 1.3% recorded in over half of the stations. This result is probably explained by environmental filtering resulting from very contrasted conditions across the different sampled systems, passive larval diffusion (Basset et al., 2007) and the randomness of the colonization process by species of marine origin (Pérez-Ruzafa et al., 2011b), a phenomenon previously observed for benthic macroinvertebrates in Italian coastal lagoons (Basset et al., 2007), fish in Atlantic and Mediterranean coastal lagoons (Pérez-Ruzafa et al., 2007b) and macrophytes in estuaries and lagoons (Pérez-Ruzafa et al., 2011c). Consequently, unique assemblages of benthic macroinvertebrates are present in each lagoon and even at smaller spatial scales (*i.e.* stations inside a lagoon), which could explain why we were able to explain only 22% of the macrobenthic community variability. This high level of lagoon-specificity present across multiple biological compartments in coastal lagoons warrants the need to go beyond a species-based approach and towards a trait-based approach (Lacoste et al., 2023), as previously outlined using macrofauna size (Basset et al., 2008; Brauko et al., 2020; Sigala et al., 2012). Such an approach could build upon the saprobity concept (Tagliapietra et al., 2012) to select relevant biological traits and ultimately help us build a more general framework around macrobenthic community assembly rules in the line of what Pérez-Ruzafa et al. (2013) did for fish by revisiting the r and K-strategies. Nonetheless, for such an approach to be relevant, we also need to increase our knowledge of the life-cycle of lagoon benthic invertebrates (Brauko et al., 2020) and to create a harmonized Mediterranean wide trait database for these organisms (Lacoste et al., 2023).

Another interesting avenue to pursue could be to investigate intra-specific biological trait and/or physiological response variability (*i.e.* oxydative stress) of species like the polychaete *Hediste diversicolor* along eutrophication gradients. Indeed, this species is known to display a great diversity of life history characteristics along with morphological, biochemical and physiological differences according to local environmental conditions (Scaps, 2002). For example, the feeding behavior and diet of this species has been shown to vary according to sewage pollution levels across English estuaries (Aberson et al., 2016). Another study has related oxydative stress of the polychaete *Nephtys cirrosa* to seasonal variations of abiotic parameters like temperature and salinity in the Ria de Aveiro (Magalhães et al., 2019). Furthermore, *H. diversicolor,* a species we found to be associated to eutrophic lagoons, is reported to present a great diversity of feeding modalities (*i.e.* predator, grazer, suspension-feeder and deposit-feeder (Faulwetter et al., 2014)), whereas the species associated to oligotrophic lagoons, like *Loripes orbiculatus,* present less diverse feeding modes (*i.e.* mainly based on sulfo-oxydizing bacteria (Roques et al., 2020)). This flexibility could allow organisms to better tolerate the fluctuating abiotic conditions that characterize small eutrophic lagoons, stressing that environmental stability is probably a strong driver of macrobenthic communities in coastal lagoons, as previously shown for macrophytes (Pérez-Ruzafa et al., 2011c). Overall, the plasticity several species display in environmentally constraining conditions stresses the difficulty to implement trait-based approaches in ultra-variable environmental systems like coastal lagoons, where realized trait modalities may differ significantly from the ones available in the literature, commonly acquired from non-transitional coastal ecosystems (Martini et al., 2021). Finally, a functional approach based on well-informed biological traits and biomass data could help overcome the identified taxonomic limitations and build a more holistic framework linking natural variability and eutrophication of coastal lagoons to macrobenthic communities but the intra-specific plasticity for many biological traits makes this approach questionable today.

## Conclusion

Coastal lagoons are transitional ecosystems naturally characterized by a strong variability in abiotic parameters like salinity and temperature. Superimposed onto this natural environmental variability are numerous anthropogenic disturbances, eutrophication being a historical one targeted by the European Water Framework Directive. By jointly analyzing water column, sediment and macrophyte parameters, our study shows that macrobenthic communities across 29 French Mediterranean coastal lagoons are primarily determined by (*i*) lagoon hydro-morphology (connection level with the sea, surface and depth) which further affects natural environmental variability in terms of salinity and temperature, and (*ii*) lagoon-scale benthic habitat diversity, as estimated through macrophyte morphotypes. Eutrophication and associated stressors like low dissolved oxygen act upon the existing communities by reducing taxonomic richness and diversity. Eutrophication appeared to act on lagoon macrobenthos through relatively direct mechanisms like toxicity of inorganic nitrogenous compounds, as well as indirectly via for example, the simplification of benthic habitats and sediment bio-geochemistry modifications. Furthermore, our study highlights the complex interconnections between lagoon hydro-morphology, natural environmental variability and eutrophication and their joint effect on benthic communities. Indeed, lagoon-sea connection modulates the susceptibility of each lagoon to eutrophication, high water temperatures promote eutrophication related phytoplankton blooms while eutrophication reduces benthic habitat heterogeneity, all of which affect benthic communities. Finally, benthic taxa richness and diversity also proved to be indicators sensitive to eutrophication in coastal lagoons while an AMBI like index based on redefined ecological groups for coastal lagoons, which can be naturally enriched in organic matter, has the potential of being a specific eutrophication indicator.

## Appendices

## Appendix 1 - Procedure used to calculate the beta diversity of macrophyte morphotypes (β_MM_) corresponding to each macrofauna station

### Step 1: Select the macrophyte stations corresponding to each macrofauna station

- If one macrofauna station is considered in a given lagoon, then all the macrophyte stations of that given lagoon are taken into account.
- If two or three macrofauna stations are considered in a given lagoon, then all the macrophyte stations attributed by the WFD protocol to the same section of the lagoon as the macrofauna station are taken into account.

### Step 2 : Build the macrophyte station by macrophyte morphotype matrix for each macrofauna station

For each macrophyte station of the dataset:

- If the macrophyte cover is below 25% or equal to 25%, a value of 1 is entered in the sediment column and 0 in all the others (line 1 of the example matrix below).
- If the cover is above 25%, the dominant or co-dominant taxa are determined based on the taxa specific cover data or the taxa specific biomass data, then these taxa are recoded into a morphotype (see Appendix 2) and finally, a value of 1 is entered in all the cells corresponding to the given morphotypes of the dominant or co-dominant taxa and a value of 0 is entered in the remaining cells (lines 2 and 3 of the example matrix below).

### Step 3 : Calculate the index value corresponding to each macrofauna station

- Calculate the Bray-Curtis dissimilarity between all the station pairs using the previously built station by macrophyte morphotype matrix (presence/absence)
- Average the Bray-Curtis dissimilarities

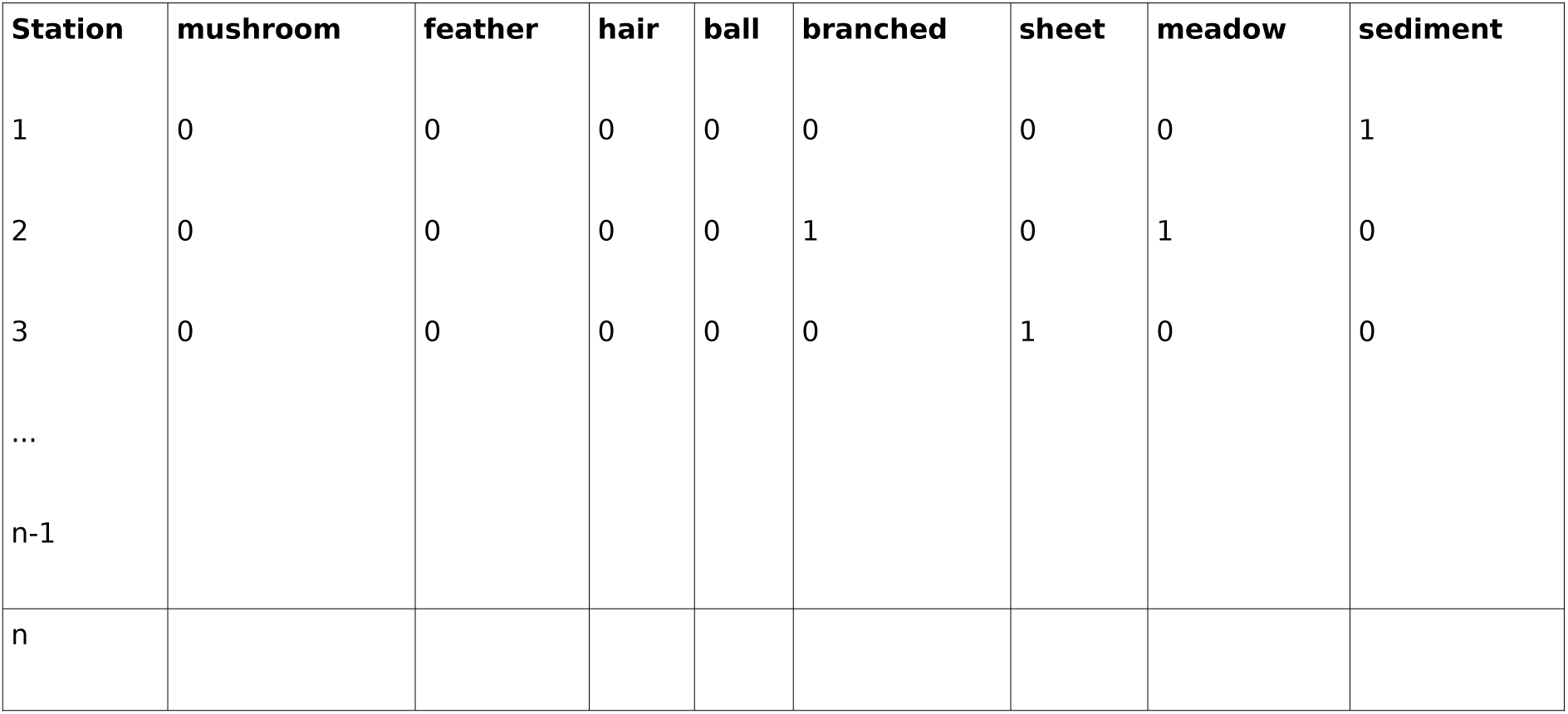
Example of a macrophyte station by macrophyte morphotype matrix used to calculate the βMM.

| Station | mushroom | feather | hair | ball | branched | sheet | meadow | sediment |
| --- | --- | --- | --- | --- | --- | --- | --- | --- |
| 1 | 0 | 0 | 0 | 0 | 0 | 0 | 0 | 1 |
| 2 | 0 | 0 | 0 | 0 | 1 | 0 | 1 | 0 |
| 3 | 0 | 0 | 0 | 0 | 0 | 1 | 0 | 0 |
| ... |  |  |  |  |  |  |  |  |
| n-1 |  |  |  |  |  |  |  |  |
| n |  |  |  |  |  |  |  |  |

## Appendix 2 - Correspondence between macrophyte taxa and morphotypes

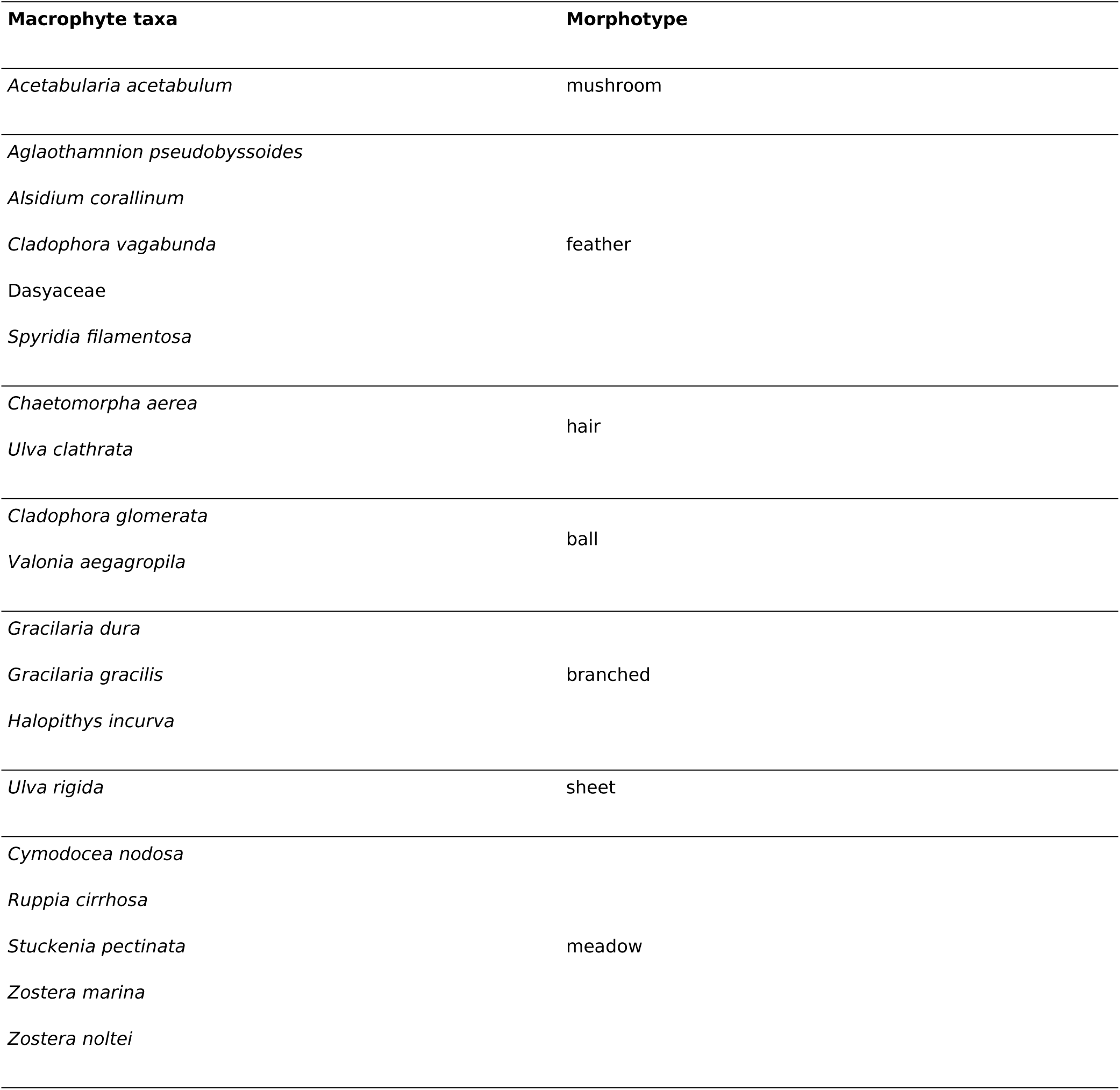

## Appendix 3

Mean ± SD of the species richness (SR), Shannon diversity (H’), inverse of Simpson dominance index (N2), macrofauna density, AMBI (AZTI’s Marine Biotic Index) and M-AMBI (Multivariate AMBI) calculated for the A and B station groups determined using a K-means partitioning based on the benthic macrofauna Hellinger-transformed mean abundances. The results of the non-parametric Kruskal-Wallis rank sum test comparing the simple and multimetric macrobenthic taxonomic indices between the A and B groups are also indicated.

|  | Group A | Group B | Kruskal-Wallis rank sum test |
| --- | --- | --- | --- |
| SR | 9.30 ± 3.48 | 17.48 ± 11.41 | $\chi^2(1) = 6.41, p = 0.011$ |
| $H'$ | 2.00 ± 0.57 | 2.36 ± 0.69 | $\chi^2(1) = 2.29, p = 0.13$ |
| $N_2$ | 3.30 ± 1.35 | 3.83 ± 1.76 | $\chi^2(1) = 0.74, p = 0.39$ |
| Macrofauna density (ind.m<sup>-2</sup>) | 2 280 ± 1 372 | 5 113 ± 5 205 | $\chi^2(1) = 0.98, p = 0.32$ |
| AMBI | 3.03 ± 0.53 | 2.87 ± 1.08 | $\chi^2(1) = 0.01, p = 0.92$ |
| M-AMBI | 0.43 ± 0.08 | 0.53 ± 0.14 | $\chi^2(1) = 5.76, p = 0.016$ |

## Appendix 4

Coordinates (biplot scores for quantitative variables and centroids for lagoon-sea connection) on the first and second axis of the redundancy analysis (RDA) plots, marginal effects and conditional effects of the 8 environmental variables selected to explain the macrobenthic community variability. After the abbreviated name of each environmental variable is indicated the type of transformation used to normalize it (log_e_: natural logarithm and sqrt: square-root). [Chla] and [DIP] is the water concentration in chlorophyll *a* (µg.L^-1^) and dissolved inorganic phosphorous (µmol.L^-1^), respectively. Temp (°C) is the lagoon water temperature. O_2_ sat (%) is the water oxygen saturation level. OMC (%) is the sediment organic matter content. CV stands for coefficient of variation.

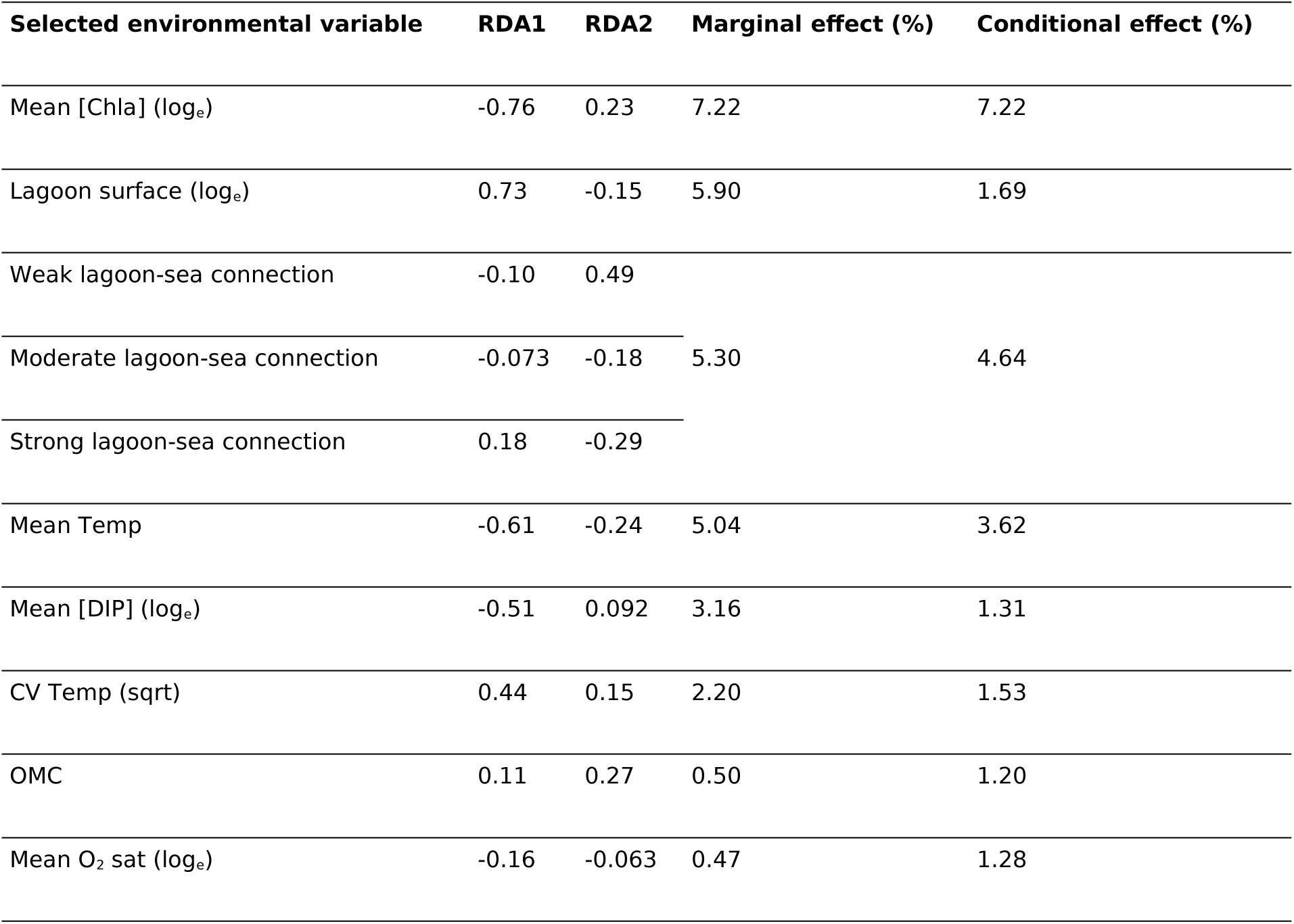

## Appendix 5

Partial redundancy analysis plots (pRDA, axis 1 and 2) focusing on (a) the benthic macrofauna sampled across 41 stations located in 29 lagoons along the French Mediterranean coast (see Table 1 for the full lagoon names and corresponding stations) as explained by keeping constant the natural environmental variability variables (lagoon-sea connection, lagoon surface, mean and CV water temperature, mean oxygen saturation and sediment organic matter content) and letting the eutrophication variables vary (mean concentration in chlorophyll *a* (mean [Chla]) and mean dissolved inorganic phosphorous (mean [DIP]), (b) the environmental variables left to vary where mean [Chla] and mean [DIP] are log_e_ transformed (see Tables 1 and 2 for the full environmental variable names and values) and (c) the macrofauna taxa, where only taxa with a goodness of fit (cumulative proportion of variance explained by the first two axis) above 0.23 are plotted. In (a), inter-station distances are euclidean while in (b) and (c) angles between all vectors reflect linear correlations. Across all plots, each station (point) is represented using a symbol for its salinity type (triangle: oligo- and meso-haline and circle: poly- and eu-haline) and a color (red: group A and blue: group B) for its group membership (K-means partitioning) based on Hellinger-transformed mean abundances.

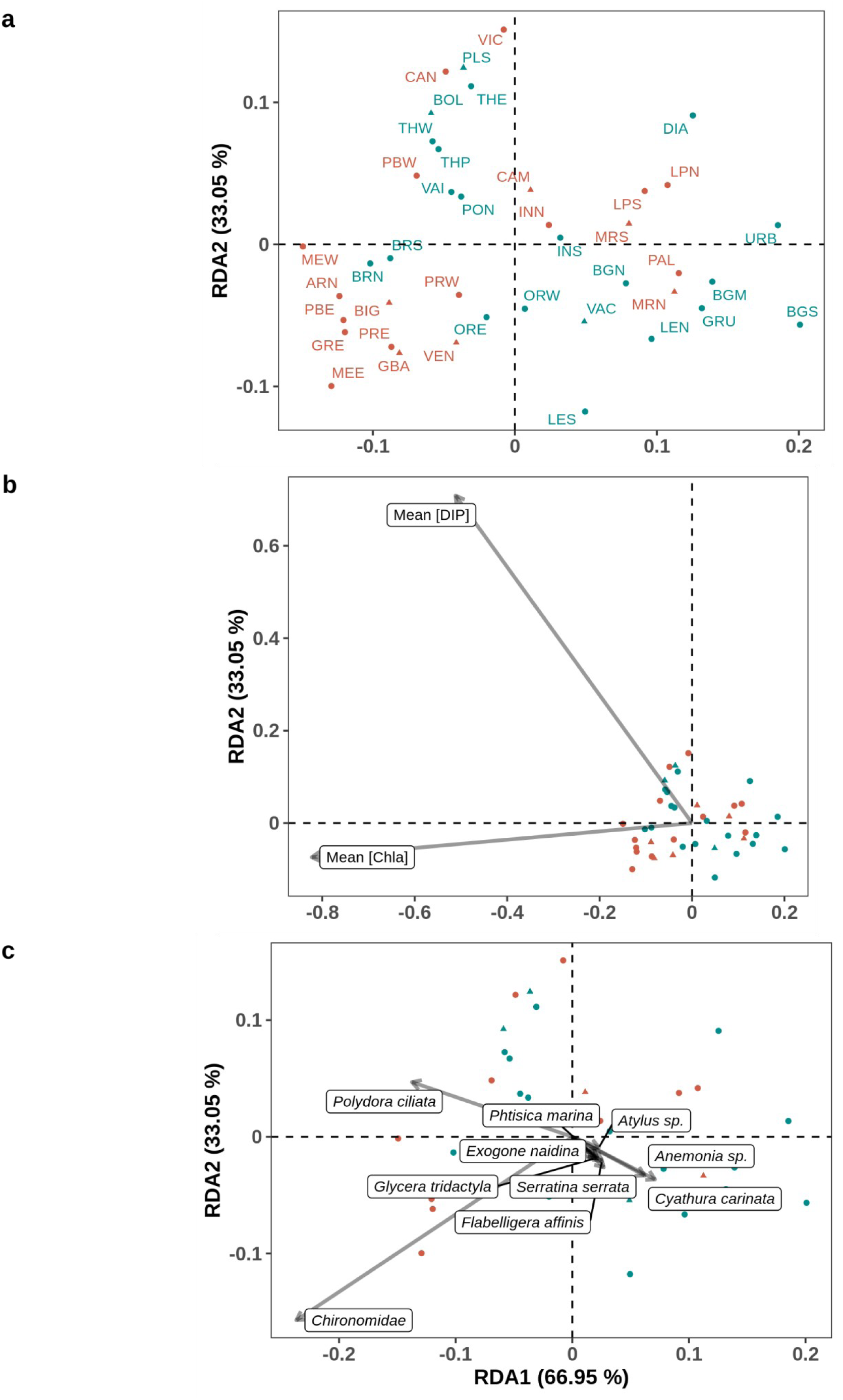

## Appendix 6

Partial effect of mean chlorophyll *a* concentration ([Chla]) on (a) species richness (SR) once the effect of mean temperature and mean salinity has been removed, (b) Shannon diversity (H’) once the effect of lagoon-sea connection, CV [TN], CV [TN]^2^ and β_MM_ has been removed and (c) M-AMBI once the effect of lagoon-sea connection and CV [NO_x_] has been removed (see Table 3 for the parsimonious linear models) across 41 stations located in 29 lagoons along the French Mediterranean coast (see Table 1 for the full lagoon names and corresponding stations). The dashed lines correspond to simple linear regression models and the shaded zone to the confidence interval. Each station (point) is represented using a symbol for its salinity type (triangle: oligo- and meso-haline and circle: poly- and eu-haline) and a color (red: group A and blue: group B) for its group membership (K-means partitioning) based on Hellinger-transformed mean abundances.

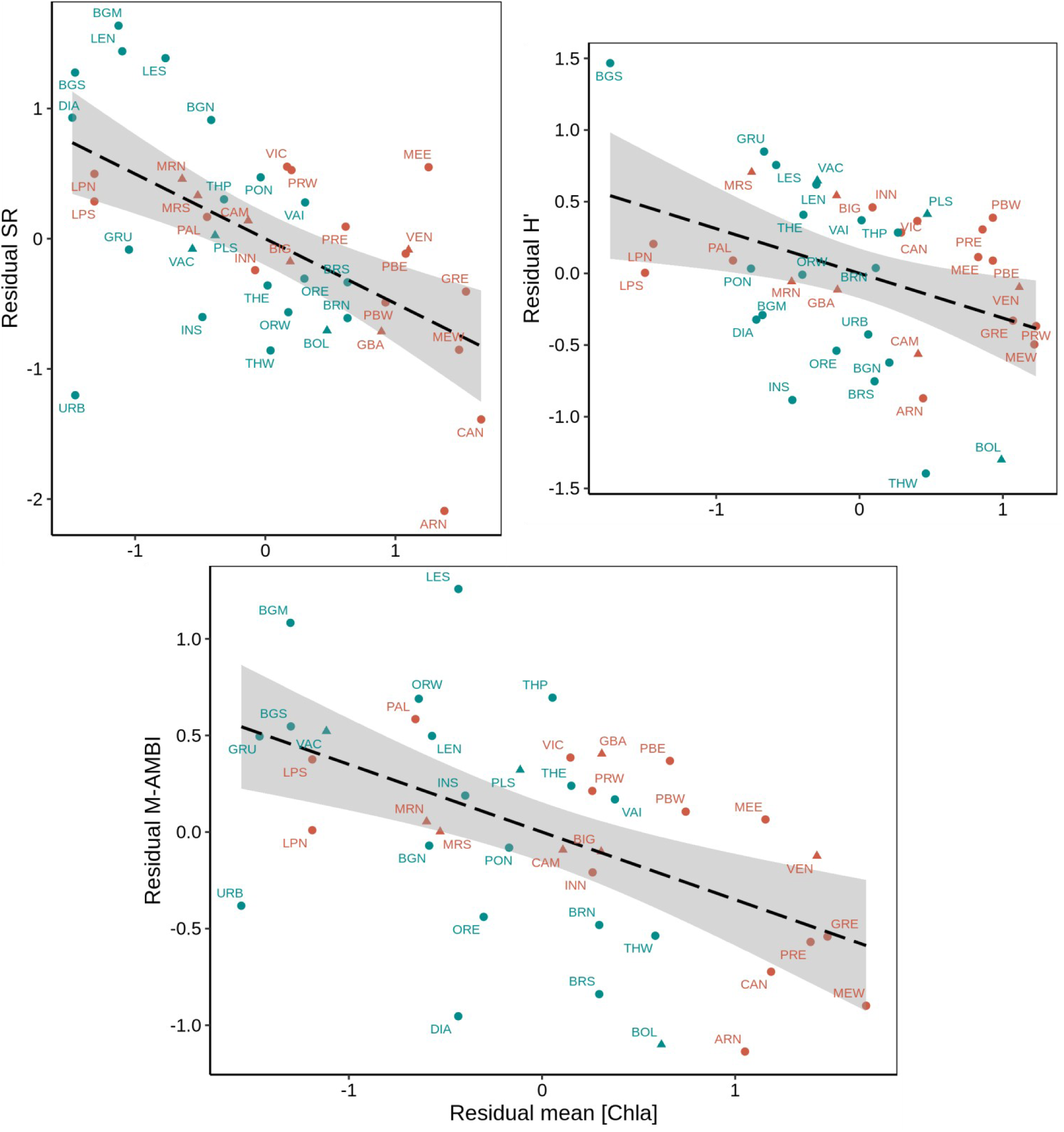

## Acknowledgements

The authors would like to thank all the members of the LERLR and LERPAC Ifremer labs as well as all the managers of the different French Mediterranean lagoons for sampling and analyzing the water column parameters, the sediment parameters and the macrofauna. A special thanks to Corinne Pélaprat (STARESO), Sébastien Thorin (CREOCEAN), Thibault Schwartz (CREOCEAN) and Daniel Martin (Centre d’Estudis Avançats de Blanes – CEAB) for sampling, identifying, and analyzing the macrofauna. The authors are also extremely grateful to Aurélie Fouveau (Ifremer, LERBN) for extracting the ecological group values, Olivier Boutron (Tour du Valat) for the information on the Vaccarès lagoon system and Marion Richard (Ifremer, LERLR) for the fruitful discussions. Finally, we would like to thank all our close colleagues from the LERLR (Ophélie Serais, Elise Bellamy, Aurélien Bouquet, Elise Caillard, Nicolas Cimiterra, Christine Felix, Elise Foucault, Camille Gianaroli, Yannick Gueguen, Elise Lacoste, Franck Lagarde, Julie Le Ray, Danièle Martin, Grégory Messian, Dominique Munaron, Hervé Violette) for creating such a cheerful, dynamic, and enthusiastic work environment, despite all the COVID-19 ups and downs.

## Data, scripts, code, and supplementary information availability

Data, scripts, and code are available online on Zenodo: https://doi.org/10.5281/zenodo.10825165

Supplementary information is available online on biorxiv as part of the preprint: https://www.biorxiv.org/content/10.1101/2022.08.18.504439

