## Supplementary material for "Disentangling the effects of eutrophication and natural variability on macrobenthic communities across French coastal lagoons"

**Supplementary material 1**

Data gathered for the Water Framework Directive (WFD) and used to estimate the missing organic matter content (OMC) values at the Arnel (ARN), La Palme North (LPN), La Palme South (LPS), Prévost West (PRW) and Vic (VIC) stations in 2006 with the calculation method used for the estimation: mean of the 3 or 4 closest stations or raw value of the closest station (see supplementary material 2 for the maps showing each lagoon and the location of the different stations).

| Macrofauna station (name used in the maps if different) | WFD closest sediment stations (distance between macrofauna and sediment station) | OMC (%) | Estimated OMC at the macrofauna station (%) and calculation | Year of the macrofauna sampling | Year of WFD sediment sampling |
| --- | --- | --- | --- | --- | --- |
| LPN (LAP) | LAP 9 (339 m) | 1.50 | 4.68 (mean) | 2006 | 2007 |
|  | LAP 10 (334 m) | 8.50 |  |  |  |
|  | LAP 11 (463 m) | 6.93 |  |  |  |
|  | LAP 12 (702 m) | 1.80 |  |  |  |
| LPS (LAPS) | LAP 4 (338 m) | 1.53 | 1.61 (mean) | 2006 | 2007 |
|  | LAP 5 (285 m) | 1.57 |  |  |  |
|  | LAP 7 (326 m) | 1.73 |  |  |  |
| ARN | ARN 3 (310 m) | 6.03 | 7.88 (mean) | 2006 | 2006 |
|  | ARN 4 (563 m) | 8.37 |  |  |  |
|  | ARN 5 (526 m) | 8.80 |  |  |  |
|  | ARN 6 (707 m) | 8.33 |  |  |  |
| PRW | PRW 2 (114 m) | 8.40 | 8.40 (raw) | 2006 | 2006 |
| VIC | VIC 12 (542 m) | 10.93 | 13.05 (mean) | 2006 | 2006 |
|  | VIC 13 (536 m) | 13.90 |  |  |  |
|  | VIC 17 (476 m) | 12.97 |  |  |  |
|  | VIC 18 (453 m) | 14.40 |  |  |  |

**Supplementary material 2**

Maps of the La Palme, Arnel, Prévost and Vic lagoons with in red, the points sampled for the WFD sediment evaluation and in black, the points sampled for the macrofauna and considered in this study.

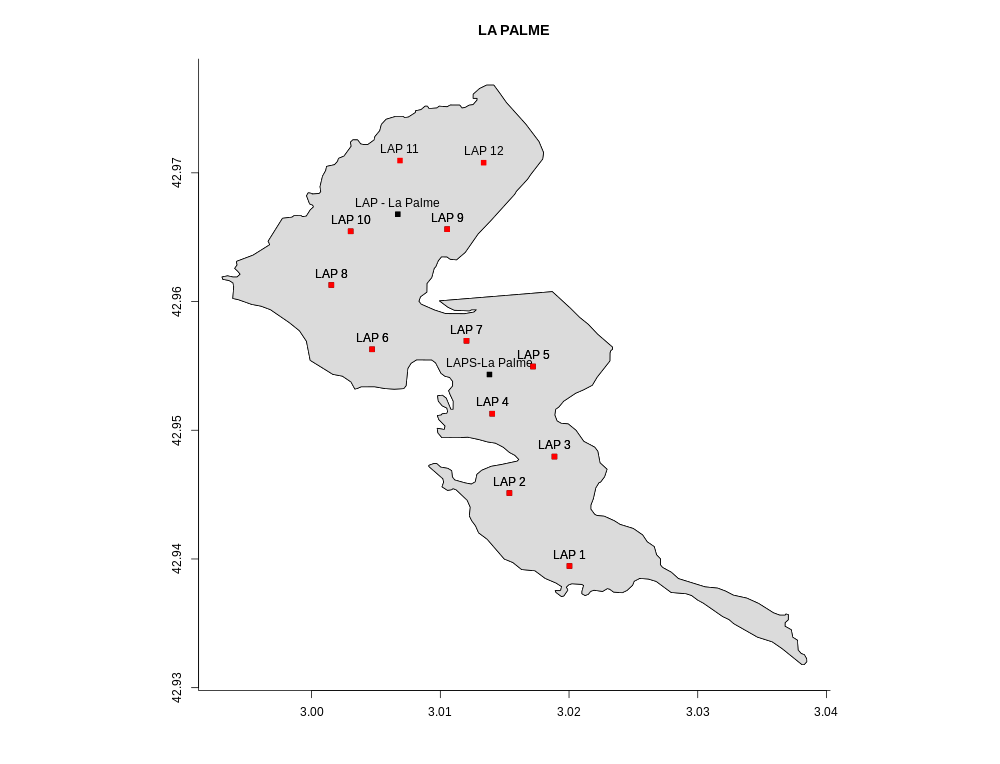

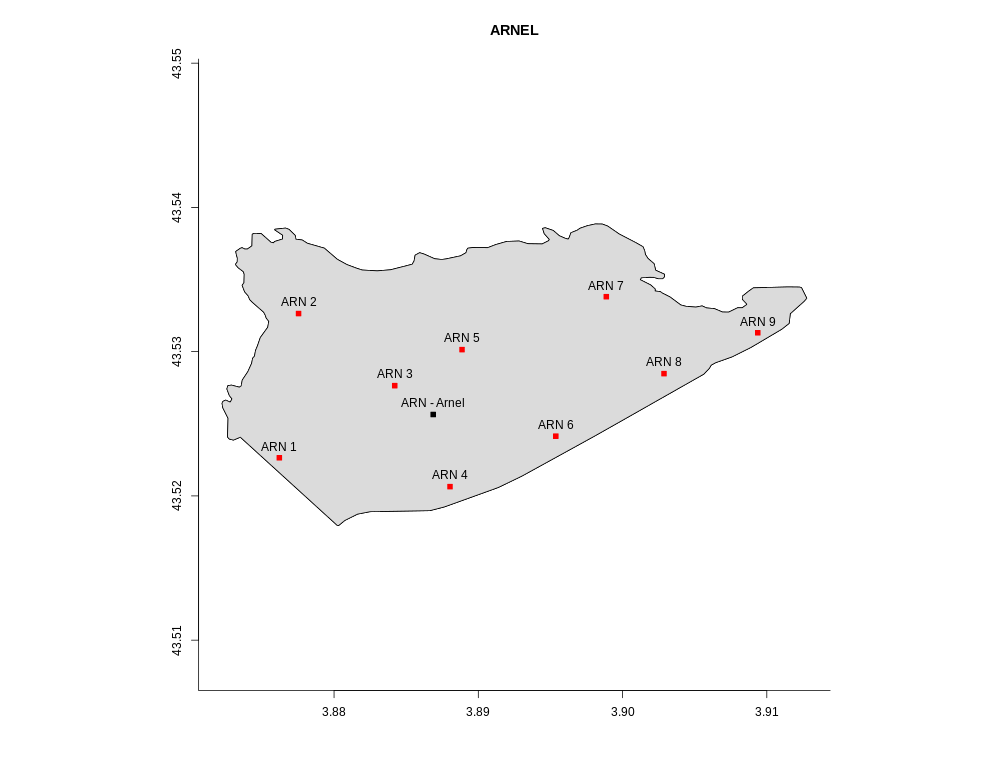

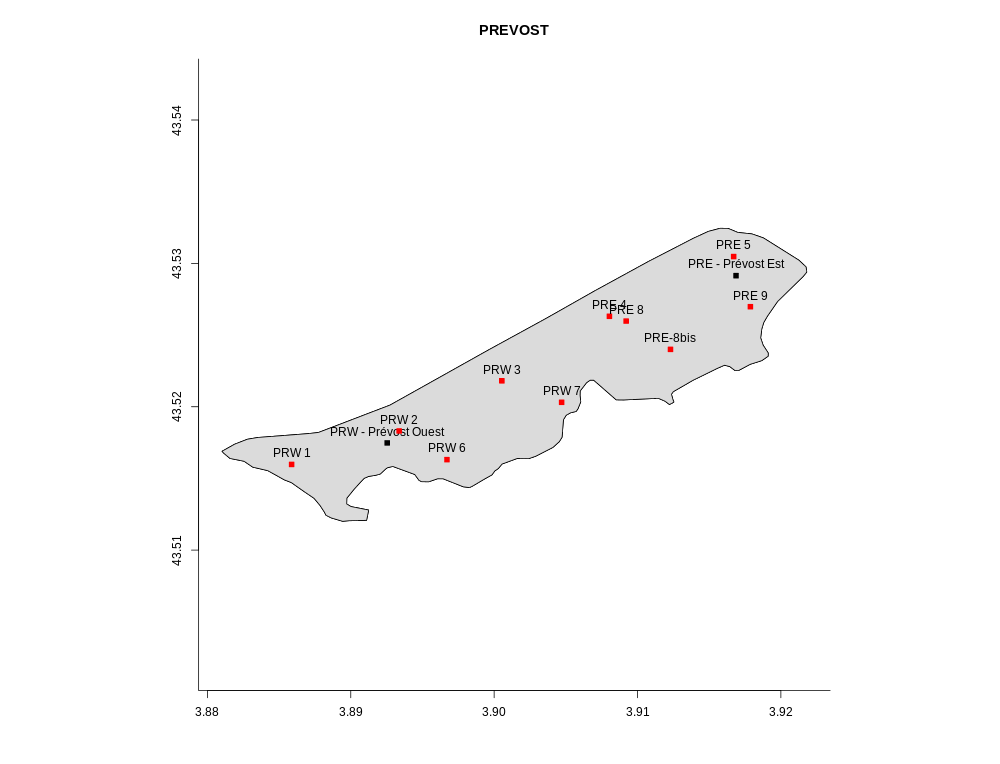

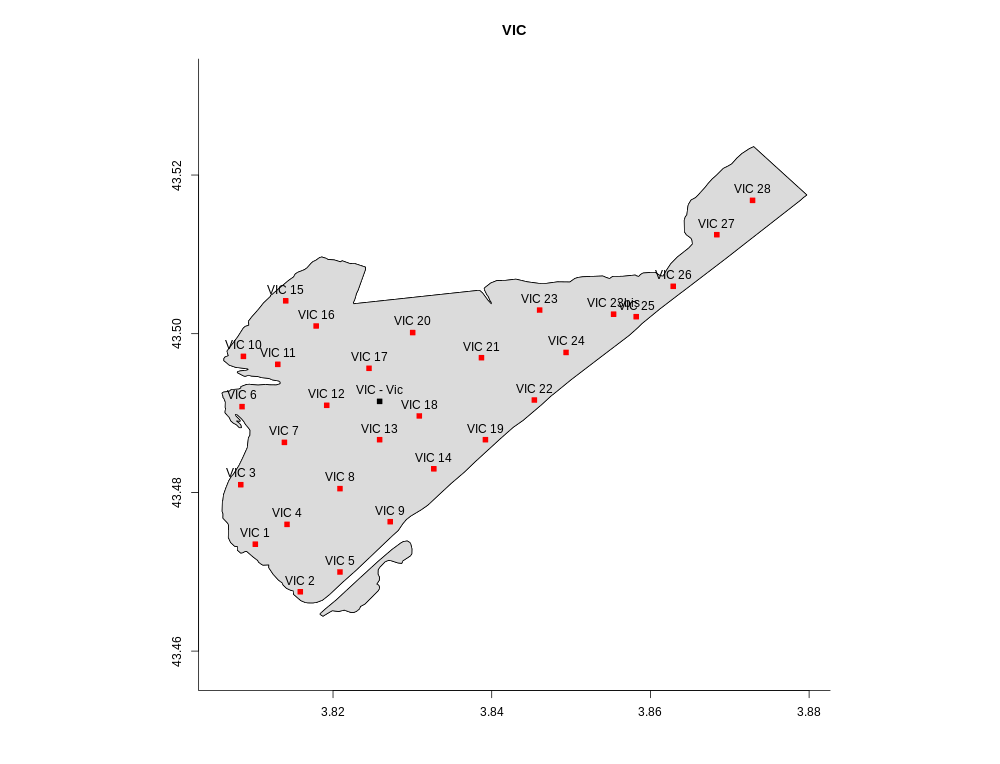

**Supplementary material 3**

Comparison of M-AMBI results on each station obtained using reference conditions of our dataset versus M-AMBI results using the reference conditions established by (i) the Water Framework directive in France (right panel, SR=46, H’=4.23 and AMBI=0.60) and using two different reference sites (ii) Leucate lagoon (center panel, SR=46, H’=3.66 and AMBI=2.11) or (iii) Thau lagoon (left panel, SR=27, H’=3.73 and AMBI=0.31) or

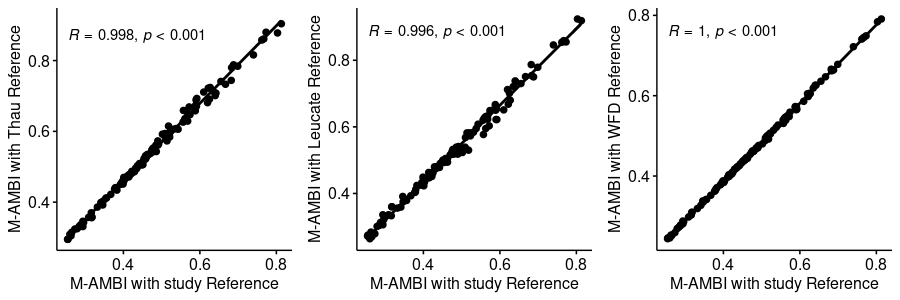

**Supplementary material 4**

Comparison of M-AMBI results on each station calculated at replicate level and then averaged at station level (our study) versus M-AMBI results directly calculated at station level (as recommended in the WFD).

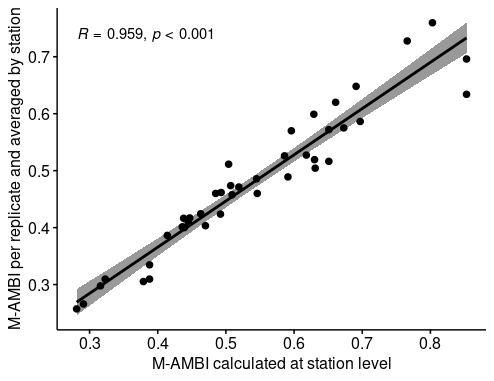
